# Influence of RNA circularity on Target RNA-Directed MicroRNA Degradation

**DOI:** 10.1101/2022.04.11.487822

**Authors:** Federico Fuchs Wightman, Jerónimo Lukin, Sebastián Giusti, Michael Soutschek, Laureano Bragado, Berta Pozzi, Paula González, Juan P. Fededa, Gerhard Schratt, Rina Fujiwara, Jeremy E. Wilusz, Damián Refojo, Manuel de la Mata

## Abstract

A subset of circular RNAs (circRNAs) and linear RNAs have been proposed to “sponge” or block microRNA activity. Additionally, certain RNAs induce microRNA destruction through the process of Target RNA-Directed MicroRNA Degradation (TDMD), but whether both linear and circular transcripts are equivalent in driving TDMD is unknown. Here we study whether circular/linear topology of endogenous and artificial RNA targets affects TDMD. Consistent with previous knowledge that *Cdr1as (ciRS-7)* circular RNA protects miR-7 from Cyrano-mediated TDMD, we demonstrate that depletion of *Cdr1as* reduces miR-7 abundance. In contrast, overexpression of an artificial linear version of *Cdr1as* drives miR-7 degradation. Using plasmids that express a circRNA with minimal co-expressed cognate linear RNA, we show differential effects on TDMD that cannot be attributed to the nucleotide sequence, as the TDMD properties of a sequence often differ between its circular and linear forms. By analysing RNA sequencing data of a neuron differentiation system, we further detect potential effects of circRNAs on microRNA stability. Our results support the view that RNA circularity influences TDMD, either enhancing or inhibiting it on specific microRNAs.

## Introduction

Circular RNAs (circRNAs) are long-known regulatory RNAs that have gained increasing attention since the first reports that highlighted their high diversity and abundance (1–4). CircRNAs are covalently closed structures that originate from pre-mRNA backsplicing and therefore lack a poly-A tail and 5’-cap. As these terminal modifications are the entry points for the microRNA (miRNA) effector machinery (5, 6), circRNAs seem largely immune to posttranscriptional degradation by miRNAs. Current knowledge points towards a diversity of both nuclear and cytoplasmic functions for different individual circRNAs, and several findings have suggested that some circRNAs regulate gene expression by working as miRNA “sponges” (7–9).

Whether circRNAs act on miRNAs by blocking their ability to bind linear mRNA targets, affecting their stability, or a combination of both remains to be determined. In particular, it is unclear whether some circRNAs are active in driving Target Directed MicroRNA Degradation (TDMD), a mechanism that has emerged as central in affecting miRNA turnover (10–14). During TDMD, targets with extensive base-pair complementarity towards the 3’ end of the miRNA –i.e. displaying no more than 3 mismatches in addition to a central bulge– lead to miRNA degradation, reversing the logic of canonical miRNA target silencing in which the target RNA is degraded. TDMD induces a conformational change on Argonaute (AGO) proteins that leads to their poly-ubiquitination and degradation, rendering the loaded miRNAs unprotected and susceptible to degradation by general nucleases (15, 16). Unlike sponging, TDMD-inducing targets act catalytically even at sub-stoichiometric levels, resulting in the most selective and potent miRNA degradation mechanism described to date. This phenomenon indeed seems to explain the half-lives of most naturally unstable miRNAs (15, 17–19). A priori, target circularity should not represent an impediment to regulating miRNAs through TDMD, and the high stability of circRNAs could even be an advantage for this activity. On the other hand, previous reports have shown that extended miRNA:target basepairing *per se* is not enough to trigger TDMD and that miRNA binding must also occur within a conformationally flexible region of the target for TDMD to be active (13, 14, 20). This property could be affected by RNA circular topology. However, to date, no circRNA has been described to drive TDMD and the only available evidence suggests that circRNAs might instead lead to miRNA stabilization (21, 22).

CircRNAs are typically coexpressed with their cognate linear RNAs from a common host gene. However, the circRNA:linear RNA ratio occurs in different proportions, with a subset of circRNAs reaching higher levels than their cognate linear isoforms (7, 8, 23). To reveal insights into circRNA-specific functions and mechanisms, a plethora of publications have relied on circRNA overexpression using various inverted repeat-containing vectors (1, 2, 24–30). Yet, the capability of plasmid-based methods to overexpress exogenous circRNAs free from overlapping, “leaky” linear RNA expression remains questionable. Thus, attributing any observed effects to the overexpressed circRNA while not rigorously controlling for the potential role of the undesired coexpressed linear transcripts represents a potential pitfall (31–33, 7, 34–36).

A key remaining question in the field is whether the circular nature of circRNAs is intrinsically or mechanistically linked to their molecular functions. In this study, we aimed to elucidate whether the linearity or circularity of targets function differently to affect miRNA stability and function. We initially examined the well described *Cdr1as (ciRS-7)*-miR-7-Cyrano network of noncoding RNAs, where the lncRNA Cyrano destabilizes miR-7-5p through TDMD (19). We confirmed that the circRNA *Cdr1as* protects miR-7-5p (21), while expression of an artificially linear version of *Cdr1as* surprisingly triggers TDMD. Along this line, we found that linear *Cdr1as* is unable to rescue the loss of the endogenous circRNA, suggesting that the circular topology of *Cdr1as* is crucial for avoiding TDMD. Using a strategy that allows the expression of an artificial circRNA with limited expression of the counterpart linear transcript, we show that circular and linear transcripts differ in their TDMD properties. Analogous to *Cdr1as*, one of the tested artificial circular RNAs failed to induce TDMD in neurons while a different artificial circRNA induced TDMD in a cell line, with their respective counterpart linear RNAs each behaving differently in both cases. Our data also present the first evidence that circRNAs are capable of effectively triggering TDMD. Moreover, through computational analysis, we propose that certain circRNA-miRNA interactions might lead to effects on miRNA stability, representing a potentially active mechanism during neuron-like differentiation.

## Materials and methods

### Plasmid construction

Unless otherwise specified, the lentiviral vectors are based on pRRLSIN.cPPT.SYN.WPRE (17).

Linear *Cdr1as* (lin*Cdr1as*) was amplified by PCR from rat genomic DNA (see primers at Supplementary table 1). Following gel purification (QIAquick Gel Extraction Kit) and cloning into pCR II-Blunt-TOPO (ThermoFisher), it was then subcloned into pRRLSIN.cPPT.SYN.WPRE. The linear *Cdr1as* version lacking the sh*Cdr1as* target site –lin*Cdr1as*^671–^ was generated by PCR amplification of a *Cdr1as* fragment lacking a 174-bp 3’-terminal segment that encompasses the miR-671 binding site, and re-cloning it into the original backbone’s BamHI-SalI sites (Supplementary table 1). The linear transcript used as a negative control consists of GFP expressed from pRRLSIN.cPPT.PGK-GFP.WPRE (Addgene plasmid 12252).

The pri-miR-132 and pri-miR-124 expressing constructs were made by amplifying the pri-miRNA fragments (Supplementary table 1) from miRNASelectTM pEGP-mmu-mirna expression vectors (Cell Biolabs) and subsequently cloning them into the BamHI-BsrGI sites of pRRLSIN.cPPT.SYN.WPRE.

The shRNAmiR (sh*Cdr1as*) is an engineered version miR-671 –previously described as a natural *Cdr1as* regulator (19, 21)– designed to be fully complementary to the circRNA and maximizing its slicing capacity (Supplementary Figure 1A). The sh*Cdr1as* lentiviral vector was constructed by replacing the pri-miRNA fragments from the previously described vectors using BamHI-NheI (Supplementary table 1), and inserting a synthetic DNA (gBlock gene fragment, IDT) by Gibson Assembly (37).

CircRNA-expressing constructs were constructed inserting gBlocks (Integrated DNA Technologies) encompassing the ZKSCAN1 upstream and downstream introns (26) flanking mCherry and bulged or seed-mutant miR-132 sites (17) downstream of a Tetracycline-inducible promoter (TREp, see Figure 2A and Supplementary table 1). Perfect or seed-mutant sites for miR-124 were subsequently inserted downstream or the circularizable region (Supplementary Figure 2A). Linear TDMD inducers expressing mCherry upstream of bulged or seed-mutant miR-132 sites were previously described (17). To replace the miR-124 sites by miR-92a sites, we introduce a g-block with the relevant sequence into the BsrGI-SalI sites using Gibson Assembly (37) (for the g-block sequence see Supplementary table 1).

Constructs expressing artificial circRNAs of different sizes are based on the pcDNA3.1(+) Laccase2 MCS Exon Vector (Addgene 69893). We inserted a perfectly matched or seed-mutant site for miR-92a using the NotI and ApaI sites located between the Laccase2 intron and the poly(A) site (see oligos in Supplementary table 1). For the long circRNA, a BamHI-NotI fragment with split-GFP circRNA from the pcDNA3.1(+) Laccase2 MCS-UNC circGFP (kindly provided by the Wilusz Lab) was inserted into the previous constructs. Finally, an insert obtained by assembly PCR with miR-218 sites (TDMD-competent or seed-mutant) was introduced into the BsrGI site located between the end of GFP and the IRES (see oligos in Supplementary table 1). The miR-218 site was based on the naturally occurring TDMD site in TRIM9, together with the 20 nucleotides flanking it (38). The linear controls were generated by introducing the corresponding PCR products into the NheI-XbaI sites of pcDNA3.1(+) (see oligos in Supplementary table 1). To generate the shorter circRNA and corresponding linear control, we amplified a shorter version of the circRNA comprising half of GFP and a part of the IRES (see oligos in Supplementary table 1) and inserted it into the AgeI-SacII or NheI-XbaI sites of the circular and linear long constructs respectively. All constructs were sequenced to verify splicing and miRNA binding sites integrity.

The lentiviral vector driving expression of FLAG/HA-AGO2 (human) from the Syn promoter (pLV-FLAG-HA_AGO2) was generated by amplifying FLAG/HA-AGO2 from pIRESneo-FLAG/HA AGO2 (Addgene plasmid 10822).

The construct for CRISPR/Cas9 genome editing of *Cdr1as* splicing sites is based on lentiCRISPRv2 (Addgene plasmid 52961), following the Zhang Lab protocol (39, 40) (Primers in Supplementary table 1).

### HEK293T culture and transient transfection

HEK293T cells were available in our institute. Cells were tested for mycoplasma contamination and only clean stocks were further used for our experiments. Cells were grown in DMEM-F12 (Gibco) supplemented with 10% (v/v) FCS and 25 U/mL Penicillin-Streptomycin (Gibco) and were plated for transfection at 100, 000 cells/well on 24-well plates. One day after plating, cells were transfected using the PEI method as previously described (17).

### Lentivirus production and transduction

Recombinant lentiviral particles were produced in HEK293T cell line. Cells were co-transfected using the PEI method with the lentiviral expression vector and two 2nd generation lentiviral packaging vectors: pMD2.G expressing the VSV-G envelope gene (Addgene plasmid 12259) and pCMVR8.74 expressing the gag/pol genes (Addgene plasmid 22036). The supernatants containing the viral particles were collected 48–72 h after transfection, concentrated using centrifugal filter units (Amicon Ultra-15, molecular weight cutoff 100 kDa, Millipore Cat. #UFC910024) as further checked as previously described (17).

### Animals used in this study

All animal tissues used in this study were obtained under experiment protocol no. No.2020-04-DR with the approval from the Comisión Institucional para el Cuidado y Uso de los Animales de Laboratorio (CICUAL) at the Instituto de Investigación en Biomedicina de Buenos Aires (IBioBA) – CONICET – Partner Institute of the Max Planck Society.

### Neuronal cultures and lentiviral transduction

Cortical and hippocampal neurons were dissected from embryonic day 16.5 and 18.5 (E16.5 and E18.5), respectively, CD1 embryos of mixed sex. Culture preparation was performed as previously described (41, 42). Briefly, cortex from CD1 mouse embryos were dissected and a neuronal suspension was prepared through Trypsin digestion and mechanical disruption of the tissue. Neurons were plated in 24 multi-well plates at a density of 80cells/mm^2^ (150.000 cells per well) and maintained in Neurobasal-A media (ThermoFisher) with 2% B27 and 0.5 mMGlutaMAX-I (ThermoFisher) at 37 °C and 5% CO2. CD1 mice for neuronal cultures were provided by our Specific Pathogen Free Animal Facility.

The euthanasia of the animals to generate primary neuronal cultures was performed under experiment protocol no. 2020-04-DR which was evaluated by the Institutional Animal Care and Use Committee of the IBioBA-CONICET according to the Principles for Biomedical Research involving animals of the Council for International Organizations for Medical Sciences and provisions stated in the Guide for the Care and Use of Laboratory Animals.

Neurons were transduced 4-7 days after plating (DIV4-7) with lentiviral constructs: lin*Cdr1as*^671-^, sh*Cdr1as*, the linear control or a combination of these, appropriately described in Results and Figures. The vectors driving each of the circRNA-expressing constructs were transduced in combination with a lentiviral construct expressing the tetracycline-controlled transactivator protein (LV-Syn-tTA). RNA was extracted at DIV11 as indicated below.

### FLAG/HA-AGO2 transfection and immunoprecipitation (AGO2 RIP)

The FLAG/HA-AGO2 expressing plasmid was transfected into HEK293T cells as described above. Immunoprecipitation of FLAG/HA-AGO2 was performed with Anti-FLAG M2 Magnetic Beads (Sigma Cat # M8823). Beads were washed twice with TBS buffer (50 mM Tris HCl, 150 mM NaCl, pH 7.4). For each immunoprecipitation (IP), one 6-cm plate with 50% confluency was used. Cells were washed once with cold PBS and lyzed in 500 μl of lysis buffer [50 mM Tris–HCl pH 7.5, 150mM NaCl, 1% (v/v) TRITON X-100, 1 mM EDTA, containing protease inhibitors (cOmplete, EDTA-free Protease Inhibitor Cocktail, Roche) and RNase inhibitor (Invitrogen)]. The lysates were incubated 30 minutes on ice, cleared by centrifugation at 16, 000 g for 10 minutes at 4 degrees and mixed with the washed beads. After 2 hours of rotation at 4 degrees, the beads were washed three times with TBS buffer. As a control for the IPs, non-transfected HEK293T cells were used. FLAG/HA-AGO2 expression and immunoprecipitation efficiency were determined by Western blot using anti-HA antibody (clone 3F10, Roche). RNA was extracted by adding Trizol reagent (Invitrogen) directly on the beads.

### Subcellular fractionation

Briefly, the circRNA-expressing construct and the linear control were transfected into HEK293T cells as described above, in 6-well plates at 50% confluency. After 48 hours, cells were harvested using 500 μl of PBS, transferred to a microcentrifuge tube and centrifuged at 500 g for 5 minutes. The supernatant was discarded, and the pellet resuspended in 350 μl of PBS plus 0.1% NP-40 (IGEPAL). 150 μl were separated and called TOTAL fraction. The remaining volume was centrifuged for 10 seconds at 10, 000 rpm. 150 μl of the supernatant were separated and called CYTOPLASM fraction. The pellet was resuspended in 150 μl of PBS plus 0.1% NP-40 (IGEPAL) and centrifuged again at 10, 000 rpm. The supernatant was discarded, and the pellet resuspended in 150 μl of PBS plus 0.1% NP-40 (IGEPAL). This was called the NUCLEAR fraction.

Out of the 150 μl of each fraction, 75 μl were used for Western Blotting and 75 μl for RNA extraction followed by reverse transcription and quantitative polymerase chain reaction (RT-qPCR).

### RNA extraction

Total RNA extractions were made using Trizol reagent (Invitrogen) following the manufacturer’s instructions.

### RT-qPCR quantification

MiRNA and *U6* expression levels were determined by using Taqman® microRNA Assays (Applied Biosystems) following the manufacturer’s instructions or using SYBR Green and step-loop RT-qPCR (see Supplementary table 3 for oligo design). MicroRNA levels were normalized to *U6* RNA levels. Standard curves for the analysed miRNAs and *U6* RNA were performed with serial dilutions of selected cDNAs, allowing calculation of relative abundances. For quantification of target mRNAs, the artificial circRNA and *Cdr1as*, total RNA was treated with RNase-free DNase I (DNA-freeTM Kit, Ambion) and reverse transcribed using random hexamers and SuperScriptTM II Reverse Transcriptase (Invitrogen) or MMLV-RT (Sigma), following the manufacturer’s instructions. The abundance of target mRNAs, artificial circRNA and *Cdr1as* were determined by SYBR green qPCR using a custom-made qPCR mix (Supplementary table 2) and specific primers (detailed at Supplementary table 3). Alternatively, INBIO highway qPCR SYBR green mix was used (Ref M130). Standard curves for the analysed amplicons were performed with serial dilutions of selected cDNAs, allowing calculation of relative abundances.

### Northern blot analysis

Northern blot analysis was performed according to standard procedures (43). A total of 10 μg RNA was loaded in each lane. The radioactively labelled probe, corresponding to the mCherry CDS fragment, was prepared using the Megaprime DNA labelling kit (Amersham Biosciences) according to manufacturer’s instructions. Alternatively, biotin-labeled oligonucleotide probes were used together with chemiluminescent nucleic acid detection kits according to manufacturer’s instructions (Thermo Scientific). The 18S RNA from the agarose gel run was used as loading control. Blots were hybridized at 65° and washed in 0.2× SSC/0.1% SDS. The blots were exposed to Phosphorimager screens and scanned with Typhoon FLA 7000 (GE Healthcare Life Sciences) or exposed to X-ray film. The relative intensities of the bands were measured by densitometry using ImageJ.

### Western blot analysis

Protein samples were separated on 12% SDS-polyacrylamide gels and transferred to PVDF membranes. Membranes were incubated with primary antibodies: anti-HA 3F10 (Rat, 1:2500), anti-Tubulin (polyclonal rabbit anti β Tubulin H-235 from Santa Cruz Biotechnology, 1:2500) and Histone-3 (polyclonal rabbit anti H3 H-0164 from Merk, 1:2500). After washing, membranes were incubated with IRDye® 800CW (LI-COR Biosciences) secondary antibodies. Bound antibody was detected an Odyssey imaging system (LI-COR Biosciences).

### Statistical analysis

R programming language was used to process information, visualize and design graphs, and perform statistical tests. Data was normalized and scaled across experiments using Unit Length Normalization. Bar, line, scatter, dot and boxplots were designed using the ggplot2 (44) and ggpubr packages. Statistical tests were done using base R and/or ggpubr (and its complement rstatix) (https://rpkgs.datanovia.com/ggpubr/index.html). For experiments where two conditions were compared, we performed unpaired t-tests, and for those where multiple comparisons were made, we used the Bonferroni correction (ns: p > 0.05, *: p <= 0.05, **: p <= 0.01, ***: p <= 0.001, ****: p <= 0.0001). For comparisons involving multiple groups we used either ANOVA followed by Tukey multiple comparisons, Generalized Linear Models (GLM) with emmeans (https://github.com/rvlenth/emmeans), or Wilcoxon rank sum test p-values (corrected with the Hochberg method for multiple comparisons) as indicated in the text. When comparing several miR-7 targets across conditions, we performed a two-way ANOVA followed by Tukey multiple comparisons, showing the adjusted p-value.

### Bioinformatic analysis of circRNA-miRNA interactions

Two publicly available datasets produced by the Chen Lab were combined to generate a comprehensive analysis taking into account miRNA, circRNA and mRNA expression during differentiation of H9 (hESC) cells to H9 forebrain neurons. From GEO Accession GSE73325 (45), we obtained data for circRNA and mRNA expression. From GEO Accessions GSE56152 and GSE63709 (46), we obtained data for microRNA expression. Noteworthy, although the same procedure for differentiation was followed (45, 46), sequencing was done on different days (D26 and D35, respectively). We realize this is suboptimal, nevertheless we proceeded with the analysis considering this the best data available to approach our questions.

Data of the interactions between circRNAs and miRNAs was retrieved from Starbase/ENCORI v3 database (http://starbase.sysu.edu.cn; (47, 48)). This particular source was selected because it combines site prediction by several programs with experimental validation (e.g., by CLIP). The main parameters chosen for downloading through their API were: assembly = hg19, geneType= circRNA & miRNA, clipExpNum=1, program= all, programNum= 2, target= all and cellType= all.

A condensed spreadsheet summarizing all the previously mentioned data can be found at Supplementary Dataset 3. The analysis was done in R, briefly:

- The expression tables were loaded and merged using the Tidyverse package (49) and/or the readxl package (50), while the circRNA-miRNA interactions table was joined using the Fuzzyjoin package (https://CRAN.R-project.org/package=fuzzyjoin) to consider genomic locations of both the circRNAs and the miRNA sites.
- Log_2_ fold changes across differentiation were calculated for each type of RNA.

Analysis from a miRNA perspective:

- An absolute number of validated miRNA-specific sites (when present) was calculated on every circRNA.
- A relative number of effective miRNA-specific sites was calculated by weighing the above number (i.e. the absolute number of miRNA-specific sites on every circRNA) to the average number of backspliced junction reads before and after differentiation as an estimate of the contribution of each circRNA.
- A total number of effective sites was calculated per miRNA (referred to as “miRNA-specific effective sites” later on) by summing all the effective miRNA-specific sites among all circRNAs.
- A miRNA-specific Pairing coefficient was defined by dividing the above number (“miRNA-specific effective sites”) by the miRNÁs average abundance pre- and post-differentiation. To simplify the analysis, we separated miRNAs in quartiles of increasing pairing coefficients and performed the downstream analysis based on them.

Analysis from a circRNA perspective:

- The sum of all (i.e. pooling all miRNAs) absolute numbers of validated miRNA sites on each circRNA (when present) was calculated.
- The “pairing capacity” index for each circRNA was calculated by multiplying the above number to the number of junction reads pre-differentiation.

All graphs were programmed and illustrated using the packages ggplot2 (44), ggrepel (https://CRAN.R-project.org/package=ggrepel) and/or viridis (https://CRAN.R-project.org/package=viridis). The network diagram (Supplementary Figure 5A) was done using IGraph package (https://igraph.org/r/). Statistical analysis of the boxplots was done and added to the graphs using ggpubr (https://rpkgs.datanovia.com/ggpubr/index.html).

For pri-miRNA analysis we used the Galaxy platform (51). Briefly, raw reads extracted from GSE73325 were trimmed using Trim Galore! (https://github.com/FelixKrueger/TrimGalore) and mapped to the human reference genome (hg38) with the RNA-STAR aligner (52). Pri-miR counts were obtained from the mapped BAM files using featureCounts (53) and the annotation file (has.gff3) retrieved from miRBase (54).

### Bioinformatic prediction of TDMD-like sites on circRNAs

To predict TDMD-like sites on circRNAs, we used the scanMiR package (55). Briefly, all reported human circRNA sequences were retrieved from circBase (56) and used as an input for the *findSeedMatches* function from scanMiR to predict TDMD-like sites for every miRNA present. The data was filtered to keep only those sites on circRNAs that had at least one junction read in the neuron-like differentiation data (see above). Finally, we calculated the proportion of miRNAs with predicted TDMD-like sites (computed as those with at least one TDMD-like site on circRNAs/linear RNAs) within each of the quartiles of increasing pairing coefficients. Fisher’s Exact tests between each quartile and the least paired quartile (“-paired”) were performed to assess enrichment. The ggpie and rstatix packages were used to generate the pie charts and the ggpubr package to do the violin plot and t-tests.

### Bioinformatic analysis of miR-7 and miR-181b/d targets

Targets for miR-7 were retrieved from TargetScan 7.1 mouse (57) while the ones for miR-181b/d from TargetScan 7.2 human (57). Raw RNA-seq counts were downloaded for *Cdr1as* KO data (21) and circCSNK1G3 KD (22), accessions GSE93130 and GSE113124, respectively. A differential expression analysis was performed using the DESeq2 package (58). Graphs were made using the results of the analysis and the packages mentioned above.

## Results

### Circularity of *Cdr1as* is necessary to protect miR-7 from TDMD

In order to study whether the circular RNA nature plays a role in TDMD, we studied the case of the *Cdr1as*-miR-7 interaction (1, 2, 59). To that end, we designed tools to manipulate endogenous *Cdr1as* levels in rodent primary neurons, a model known to display high TDMD levels (60, 17, 19). A lentiviral vector was engineered to express a shRNAmiR (sh*Cdr1as*) against *Cdr1as* (Supplementary Figure 1A) that was then transduced into primary neurons at high efficiency. *Cdr1as* was effectively knocked down (Figure 1A, Supplementary Figure 1B) and miR-7 levels were reduced (Figure 1B), which is in line with previous evidence (21). *Cdr1as*, therefore, does not induce TDMD but instead helps stabilize this miRNA.

**Figure 1.**
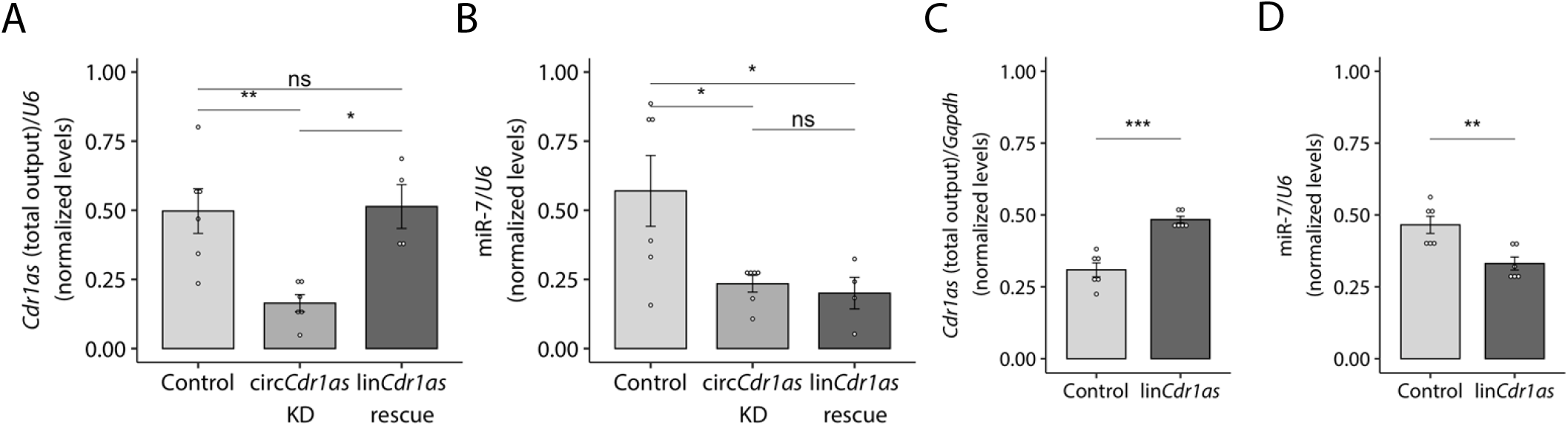
The circular topology of *Cdr1*as determines the outcome of its effect on miR-7’s stability. A *Cdr1*as total output levels (linear plus circular) measured by RT-qPCR upon transduction of either a scrambled shRNA or sh*Cdr1*as alone, or sh*Cdr1*as rescued with a linear version of *Cdr1*as (lin*Cdr1*as) lacking the *shCdr1*as site, in cortical primary neurons. n = 6 culture wells (from 3 independent primary cultures) for control and cir*CDR1*as KD; n = 4 culture wells (from 2 independent primary cultures) for lin*Cdr1*as rescue. B MiR-7 abundance measured by Taqman RT-qPCR in the same samples as in A. n = 6 culture wells (from 3 independent primary cultures) for control and cir*CDR1*as KD; n = 4 culture wells (from 2 independent primary cultures) for lin*Cdr1*as rescue. C *Cdr1*as total output levels measured by RT-qPCR upon over-expression of linear *Cdr1*as (lin*Cdr1*as) in primary hippocampal neurons. Control corresponds to a non-related (GFP-expressing) linear transcript. n = 6 culture wells (from 2 independent primary cultures) for each condition. D MiR-7 abundance measured by Taqman RT-qPCR in the same samples as in C. n = 6 culture wells (from 2 independent primary cultures) for each condition. Data are presented as mean ± SEM. Statistical significance was determined by unpaired Student’s t tests (ns: p > 0.05, *: p <= 0.05, **: p <= 0.01, ***: p <= 0.001, ****: p <= 0.0001).

To determine whether the observed effect was dependent on the circular topology of *Cdr1as*, we attempted to rescue *Cdr1as* knockdown with the expression of an artificial linear version of *Cdr1as* –lin*Cdr1as*– lacking the sh*Cdr1as* target site. Interestingly, the decrease in miR-7 abundance caused by the knockdown of endogenous *Cdr1as* could be neither rescued nor significantly enhanced by co-expressing the linear *Cdr1as*, even though the artificial lin*Cdr1as* reached expression levels similar to those of endogenous *Cdr1as* in control cells (Figure 1A-B). Remarkably, expressing lin*Cdr1as* alone –without knocking down endogenous *Cdr1as*– caused a significant destabilization of miR-7 (Figure 1C-D), consistent with TDMD being driven by the RNA expressed in an artificial linear topology. Importantly, the linear lin*Cdr1as* did not induce significant variation in the abundance of endogenous circular *Cdr1as* (Supplementary Figure 1C). To exclude the possibility that miR-7 downregulation was a consequence of changes in its transcription and/or maturation upon transduction of lin*Cdr1as*, we measured the abundance of its primary transcript (pri-miR-7a-1), passenger strands (miR-7a-1-3p and miR-7a-2-3p) and two (2) additional control miRNAs (miR-9 and -132). Our results confirmed that the guide strand miR-7-5p, but neither the primary transcript/passenger strands nor the other control miRNAs, undergo degradation upon introduction of the linear version of *Cdr1*as (Figure 1D, Supplementary Figure 1D). This result also confirms that the reduction in miR-7 is a post-transcriptional effect occurring after miRNA processing and is therefore compatible with active TDMD triggered by the lin*Cdr1*as construct (Supplementary Figure 1D). Careful inspection of the miR-7 sites present in *Cdr1as* showed that some of them indeed exhibit base pairing complementarity compatible with a TDMD-competent architecture (Supplementary Figure 1E) (14).

As an orthogonal strategy to reduce *Cdr1as* levels and rule out potential off-target or indirect effects caused by the sh*Cdr1as*, we used CRISPR/Cas9 genome editing to mutate the splicing sites of the endogenous *Cdr1as* gene (Supplementary Figure 1F). Despite an overall lower efficacy in *Cdr1as* knockdown compared to the sh*Cdr1as*, we observed a similar effect on miR-7 levels, consistent with *Cdr1as* being unable to induce TDMD on miR-7 and instead leading to its stabilization (Supplementary Figure 1G-H).

Overall, our results show that endogenous *Cdr1as* is unable to trigger TDMD on miR-7 but rather stabilizes this miRNA. Nonetheless, when expressed as an artificially linear RNA, it can engage in miR-7 degradation through TDMD, supporting the notion that the natural circular/linear topology, and not just the nucleotide sequence of a RNA target, is a crucial determinant for engaging in such type of regulation.

### Addition of microRNA perfect target sites in flanking introns improves the specificity of artificial circRNA expression from expression vectors

In order to study whether the differences in efficacy of circular and linear RNAs on TDMD are a general phenomenon, we sought to set up a system capable of expressing either artificial linear transcripts or circRNAs of identical primary sequences (Figure 2A). Both the linear and circular RNAs contain binding sites against candidate miRNAs with TDMD-competent sequence complementarity or seed-mutant controls (Supplementary Figure 2A). To express the circRNA, we flanked the region to be circularized with splice sites and flanking introns from a naturally circularized RNA (human ZKSCAN1 exons 2 and 3) that contain Alu reverse complementary sequences (RCS) that drive backsplicing (Figure 2A) (26). In order to preferentially enrich the expression of the circular over the linear variant in neurons, we further introduced perfectly matched sites against a highly expressed neuron-specific miRNA (miR-124) in the downstream intron that is lost after production of the backspliced circular product (Figure 2A). These miRNA sites remain in the linear RNA isoform, thus rendering the linear –but not the circular– RNA form susceptible to AGO2 slicing. In parallel, we expressed a linear transcript with identical sequence to the circRNA-expressing construct, but lacking the splice sites and flanking introns that induce circularization (26) (Figures 2A and Supplementary Figure 2A). The latter constructs have proved effective in triggering TDMD in primary neurons (17). All constructs were packed into lentiviral vectors and used to transduce mouse primary neurons.

**Figure 2.**
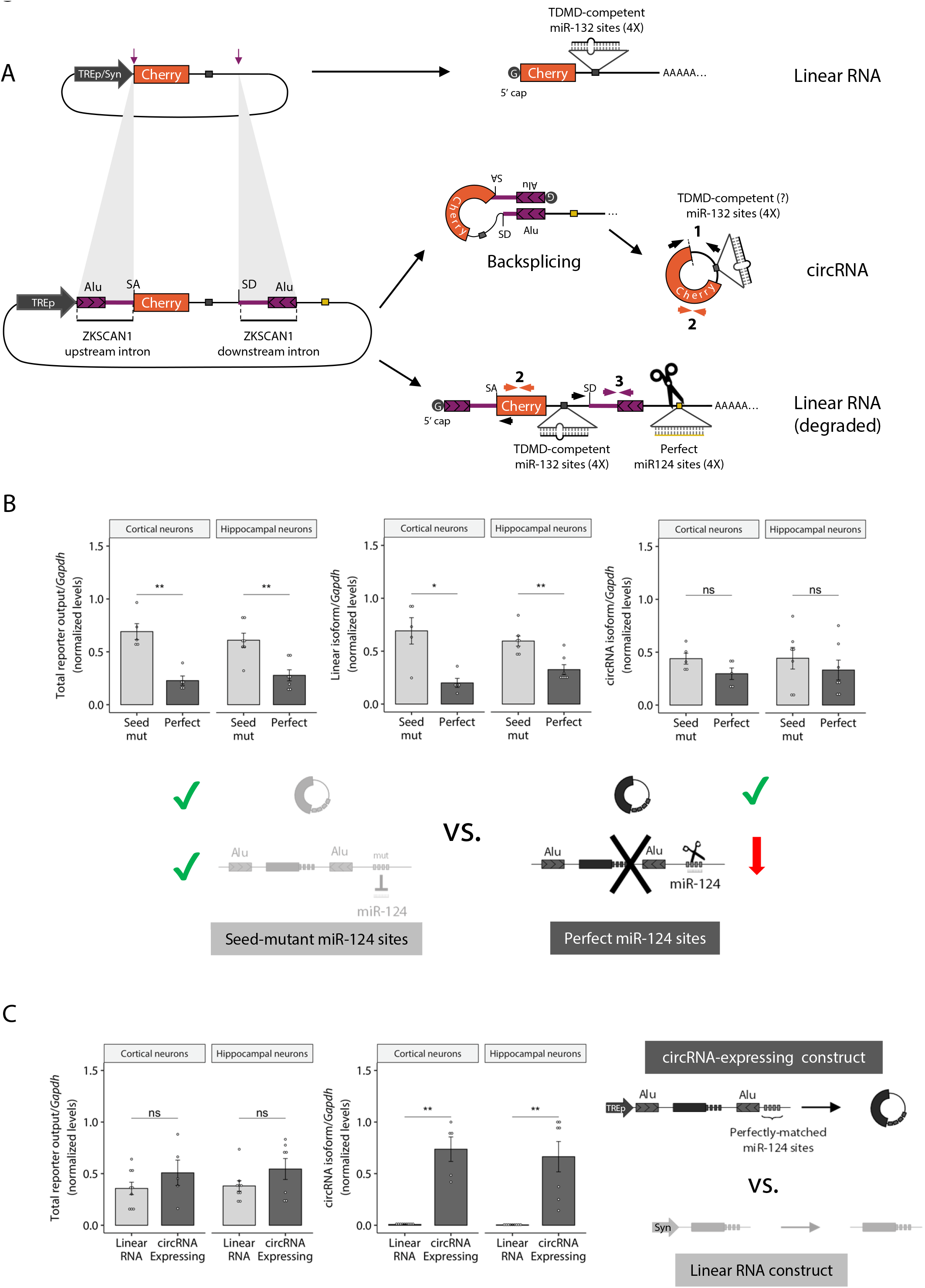
System to artificially express circRNAs reducing their overlapping, cognate linear RNA expression. A. Top: Illustration of the linear RNA expressing construct used as a positive control for TDMD (TDMD inducer). Bottom: Illustration of the circRNA-expressing construct; depicted with coloured arrows are the sets of primers used to measure the different transcript variants (circular [1], Total Output-TO [2] and linear [3]). B. RT-qPCR measuring Total output (TO, primer pair #2), linear (primer pair #3) and circular (divergent primer pair #1) RNA levels upon expression of the circRNA-expressing constructs from the tetracycline-inducible promoter (TREp) bearing perfectly matched or seed-mutant miR-124 sites for selective linear RNA degradation (see Supplementary Figure 2A). Primers are depicted in Figure 2A. n = 5 culture wells (from 3 independent primary cultures) of cortical neurons for each condition; and n = 7 culture wells (from 4 independent primary cultures) of hippocampal neurons for each condition. C. Total reporter output (left) and circRNA levels (right) upon expression of the circRNA-expressing construct or the linear RNA construct (TDMD inducer). The constructs were expressed from the tetracycline-inducible promoter (TREp) and the synapsin (Syn) promoter respectively in order to achieve similar total output levels for both constructs n = 9 culture wells (from 3 independent primary cultures) of cortical neurons for each condition; n = 9 culture wells (from 4 independent primary cultures) of hippocampal neurons for each condition. Missing points are failed culture wells/RT-qPCR reactions. In (B–C), data are presented as mean ± SEM. Statistical significance was determined by unpaired Student’s *t* tests (ns: p > 0.05, *: p <= 0.05, **: p <= 0.01, ***: p <= 0.001, ****: p <= 0.0001).

In order to validate that the constructs expressed a single circRNA species with covalently linked ends, we performed Northern blot analysis of RNase R treated or untreated samples from HEK293T transfected cells –a cell line where miR-124 is not endogenously expressed (Supplementary Figure 2B). We observed that circRNA-expressing constructs bearing either perfect or seed-mutant sites for miR-124 produced two bands: an RNase R-resistant circRNA and an RNase R-sensitive linear RNA form. This result confirms the circular topology for the artificially expressed circRNA (Supplementary Figure 2B and D). Importantly, quantification of northern blot bands correlated well with RT-qPCR measurements using primers that detected the circular isoform exclusively (Figure 2A, primer pair #1) or both the linear and circular isoforms combined –hereafter referred to as Total Output (TO)– (Figure 2A, primer pair #2), further validating the latter method for subsequent analysis (Supplementary Figure 2C). We confirmed usage of the appropriate back-splicing junction with two reverse transcriptase enzymes (MMLV-RT and Superscript II), thereby further ruling out the likelihood of artifacts due to template switching during cDNA synthesis (Supplementary Figure 2E).

To validate that our strategy was successful at expressing a circRNA while selectively degrading its counterpart linear transcript by means of miR-124, we transduced the circRNA-expressing constructs into mouse primary neurons. We observed a potent (3-4 fold) degradation of “leak” linear RNA without affecting circRNA abundance (Figure 2B). Furthermore, circRNAs were produced exclusively from the circRNA-expressing construct and not from the linear RNA-expressing construct (TDMD inducer), while both constructs produced equivalent total output levels (Figure 2C). A similar result was observed in HEK293T cells when the miR-124 sites were replaced with a perfect match against miR-92a, a highly expressed miRNA in this cell line. Under these conditions, a 2-fold reduction of the linear RNA was observed without affecting circRNA levels (Supplementary Figure 2F). Altogether, these results confirm that our system is effective in expressing circRNAs while reducing the abundance of their cognate linear RNAs, making it a generally useful tool in experiments aimed at dissecting circRNA function.

### Artificial circularization of RNAs influences TDMD efficacy

Using the constructs described above, we next explored whether linear and circular target topologies differently affect miRNA stability through TDMD in neurons, a cell type with potent TDMD activity (17). To this end, we transduced primary neurons with either linear or circRNA expression constructs bearing TDMD-competent (bulged) or seed-mutant binding sites against miR-132. The linear TDMD inducer was capable of effectively destabilizing miR-132, but the circRNA-expressing construct showed no effect on miRNA stability (Figure 3A) even when both constructs were expressed at similar overall levels (Figure 2C). Based on the performance of the circRNA-expressing construct in neurons (i.e. 3-4 fold specific reduction of linear RNA by-products and similar total transcript levels), we estimate that circRNA levels account for up to 30-50% of the total RNA expressed (compared to the ca. 25% in HEK293T cells with no endogenous miR-124). With this technical limitation in mind, we conclude that the artificial circRNA appears unable to contribute to TDMD and may even antagonize the TDMD effect driven by its co-expressed linear counterpart (Figures 2C and 3A). We confirmed that miR-132 downregulation by the linear TDMD inducer was not a consequence of changes in transcription and/or maturation of the targeted miRNA by showing that the abundance of the primary transcript (pri-miR-132) and passenger strand (miR-132-5p) of miR-132 remained unchanged (Supplementary Figure 3A). The reduction of the miR-132 guide strand was thus a bona fide post-transcriptional effect occurring after miR-132 processing and is therefore consistent with TDMD. We further examined the specificity of the linear TDMD inducer and confirmed that the levels of four additional unrelated mature miRNAs (miR-124, -128, -138 and -409) were not significantly affected (Supplementary Figure 3A).

**Figure 3.**
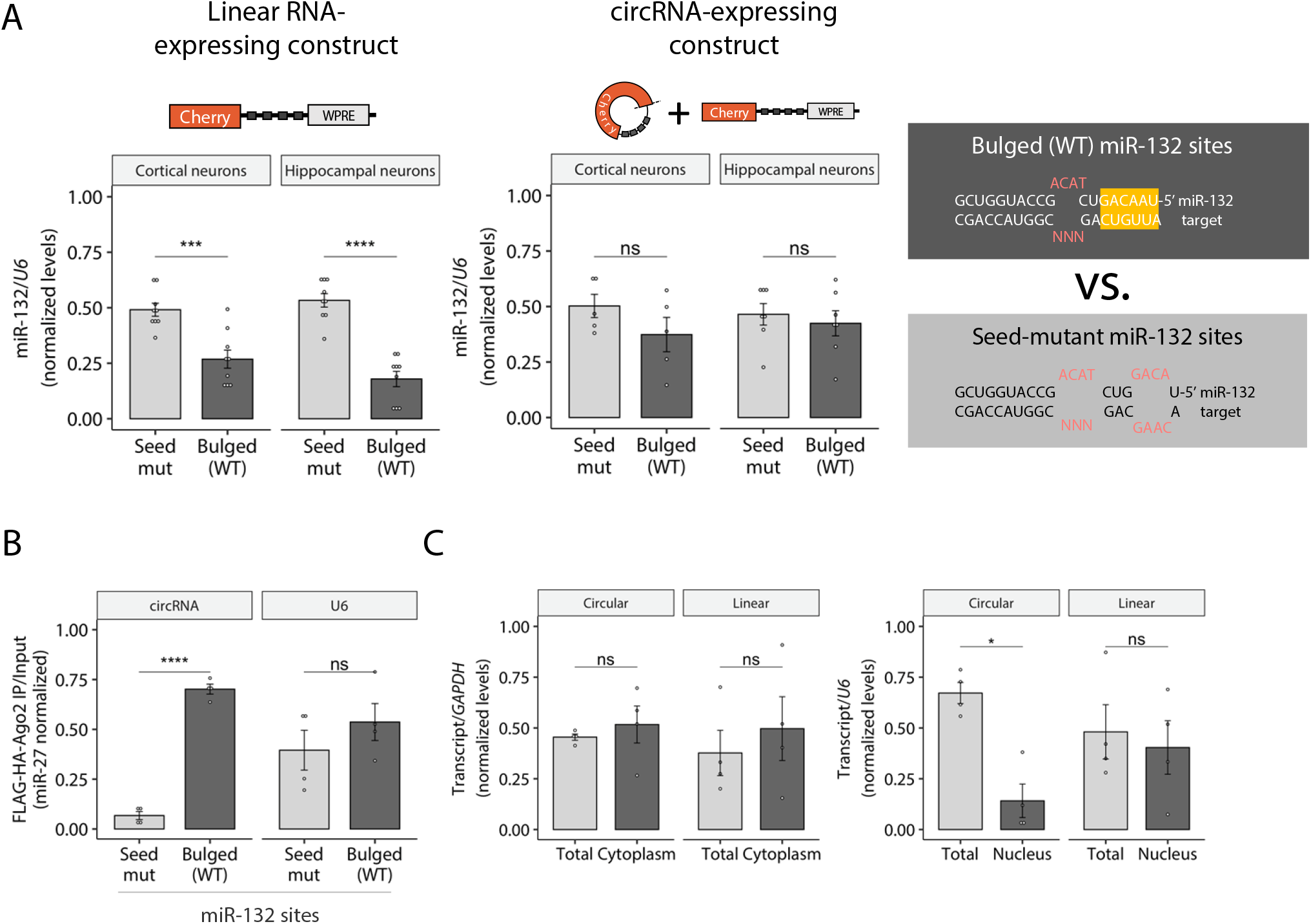
Artificial circRNA-expressing constructs are unable to trigger TDMD in primary neurons. A RT-qPCR Taqman assay showing miR-132 abundance upon transduction of the linear control (left) or the circRNA-expressing construct (right) carrying bulged (TDMD-compatible) or seed-mutant miR-132 sites. n = 9 culture wells (from 3 independent primary cultures) of cortical neurons for each condition; n = 9 culture wells (from 4 independent primary cultures) of hippocampal neurons for each condition. Missing points are failed culture wells/RT-qPCR reactions. B AGO2-Flag immunoprecipitation (RIP) followed by RT-qPCR in HEK293T cells. MiR-27a was used to normalize expression. Relative circRNA abundance was measured using circRNA-backspliced-junction specific divergent primers and normalized to miR-27a levels as an unrelated RISC-loaded miRNA not expected to be affected by the circRNA. Levels of non-specific *U6* background binding to Ago2 are shown. As an IP quality control, FLAG/HA-AGO2 input levels were shown to be similar across transfected conditions and efficiently pulled-down using anti-FLAG beads (Supplementary Figure 3B). Accordingly, miR-27, but not *U6* RNA, was efficiently co-immunoprecipitated, showing an even recovery of Ago-bound RNA across transfected conditions (Supplementary Figure 3C). n = 4 culture wells (from 2 experiments) of HEK293T cells for each condition. C Subcellular fractionation showing total vs. cytoplasmic (left) and total vs. nuclear (right) fractions, followed by RT-qPCR of the circRNA and linear RNA isoforms, normalized by *GAPDH* (for cytoplasm) or *U6* (for nucleus). Fractionation efficiency was assessed via Western Blot and RT-qPCR (Supplementary Figure 3D-E). n = 4 culture wells (from 2 experiments) of HEK293T cells for each condition. In (A–C), data are presented as mean ± SEM. Statistical significance was determined by unpaired Student’s t tests (ns: p > 0.05, *: p <= 0.05, **: p <= 0.01, ***: p <= 0.001, ****: p <= 0.0001).

Since the lack of TDMD could be attributed to the inability of the circular RNA to bind to the RISC complex, we performed an RNA immunoprecipitation experiment (RIP) by specifically pulling-down AGO2 and isolating all copurifying RNA species. We co-transfected HEK293T cells with FLAG/HA-AGO2, the different circRNA-expressing constructs and a vector for pri-miR-132 (otherwise absent in HEK293T cells), followed by anti-FLAG RIP and RT-qPCR analysis. The artificial circRNA was effectively pulled down with AGO2 when carrying bulged sites for miR-132, but not when the sites were mutated at the miRNA seed-binding region (Figures 3B and Supplementary Figure B-C). This thus demonstrated that the circular RNA is indeed able to specifically bind to the RISC complex.

We next reasoned that the observed differences could be the trivial consequence of differential localization of the circRNA relative to the linear isoform. To exclude this possibility, we performed subcellular fractionation and observed that the artificial circRNA accumulated in the cytoplasm at similar proportions relative to the linear control, with the former being only slightly lower in the nuclear compartment (Figures 3C and Supplementary Figure D-E). We conclude that the inability of the circRNA to trigger TDMD is not related to it being retained in the nucleus.

The behaviour observed for the circRNA tested up to this point led us to ask whether avoiding TDMD was a general phenomenon. To answer this, we generated additional constructs that express circRNAs from another backbone based on *Drosophila laccase2* introns which have been previously shown to produce high levels of artificial circRNAs (27). Because circRNA size could affect its structural conformation and this in turn affect both backsplicing efficiency and TDMD properties, we generated vectors expressing circRNAs of different sizes (Figure 4A). Following a similar approach as described above (Figure 2A), we introduced a perfectly matched site in the downstream intron against a highly expressed miRNA in HEK293T cells (miR-92a) in order to selectively degrade the linear RNA isoform that can be co-expressed from the vector. In parallel, we also generated the corresponding linear RNA-expressing controls. Within both the circularized region and linear RNAs, we introduced TDMD-competent (bulged) or seed-mutant binding sites against miR-218, a miRNA of low/intermediate endogenous expression in HEK293T cells. Due to the presence of a segment from the poly A signal, the linear RNAs are expected to be slightly longer than their circular equivalents.

**Figure 4.**
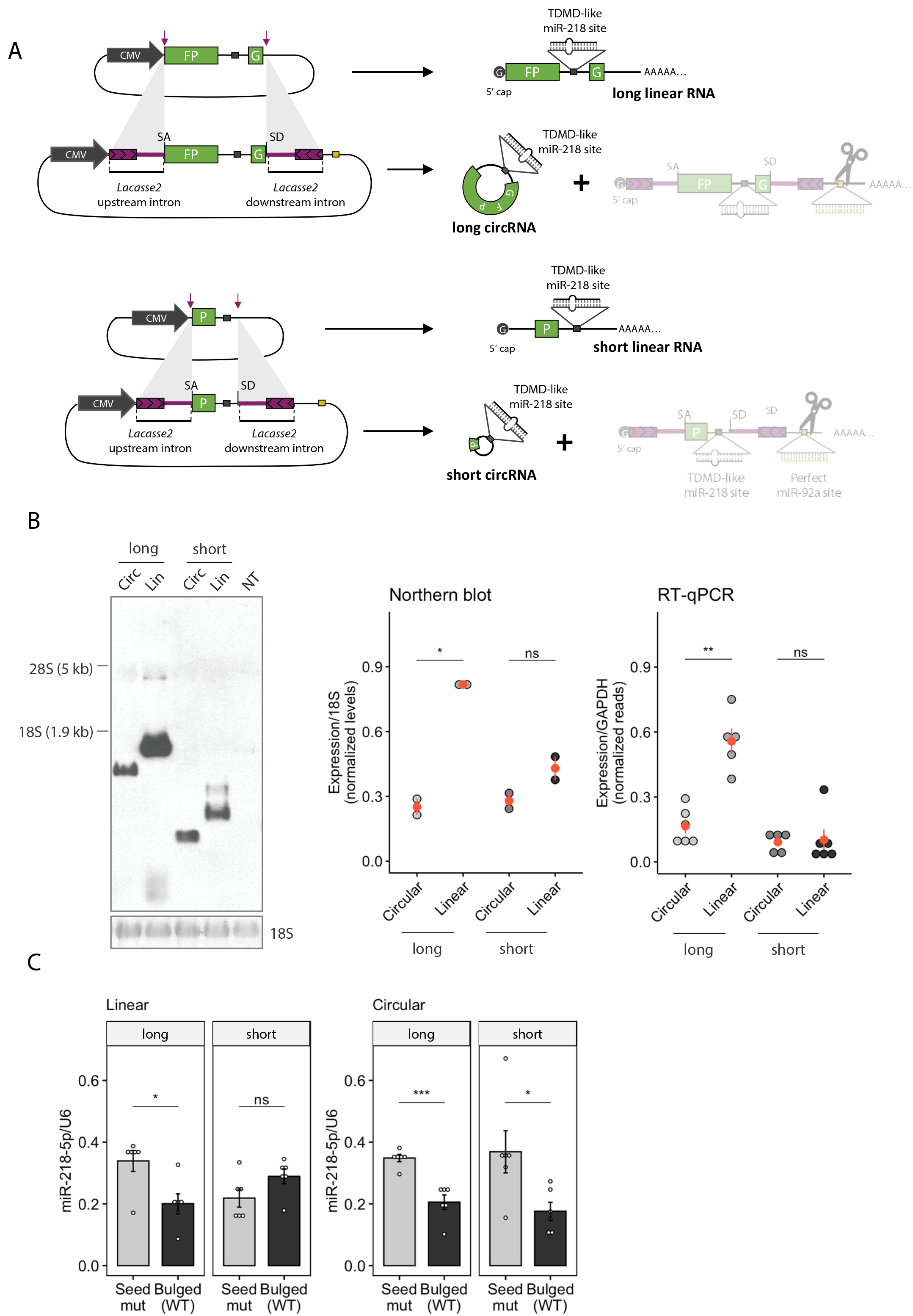
Artificial circRNA-expressing constructs trigger TDMD in HEK293T cells. A Illustration of the constructs expressing linear or circular RNA of different sizes from a backbone containing *Drosophila laccase2* introns (27). B Left: Northern blot analysis of samples from HEK293T cells expressing the constructs of the indicated length. n = 6 culture wells (from 2 independent experiments) of HEK293T cells for each condition. Right: Quantification by different methods of the linear and circular isoforms from samples of HEK293T cells expressing the indicated constructs. In red, the mean is indicated. NT are non-transfected cells. C MiR-218-5p abundance upon transduction of the linear controls (left) or the circRNA-expressing constructs (right) carrying bulged (TDMD-compatible) or seed-mutant miR-218-5 sites. n = 6 culture wells (from 2 independent experiments) of HEK293T cells for each condition. Data are presented as mean ± SEM. Statistical significance was determined by unpaired Student’s t tests (ns: p > 0.05, *: p <= 0.05, **: p <= 0.01, ***: p <= 0.001, ****: p <= 0.0001).

When transfected into HEK293T cells, the respective constructs conveniently produced essentially only circular or linear RNAs (Figures 4B and Supplementary Figure 4A). The different gel migration patterns observed were consistent with differences in both linear/circular target topology and size, with the linear RNA consisting of a slightly longer sequence due to the polyadenylation signal and the poly-A tail. Remarkably, while both circular and linear long RNAs (1408 and ca. 1570 nt respectively) induced miR-218 degradation, only the circular –but not the linear– short RNA (367 and ca. 480 nt respectively) induced TDMD (Figure 4C). Importantly, neither construct significantly affected the primary transcripts of miR-218 (pri-miR-218-1 and -2) nor an unrelated miRNA (miR-9-5p), consistent with a post-transcriptional TDMD effect of the construct acting on miR-218 (Supplementary Figure 4B). This result was in clear contrast to the previous one shown in neurons, where only the linear RNA –but not the circular RNA– induced TDMD (Figure 3).

Taken together, our results show that linear and circular RNAs counterparts can differ in their TDMD properties, supporting the idea that target RNA linear/circular topology is an important factor in determining the efficacy of TDMD.

### CircRNAs potentially affect the stability of dozens of microRNAs across neuron-like differentiation

Based on our results, we hypothesized that circRNAs might possess the ability to influence miRNA stability by either driving or evading TDMD. Yet, whether this type of regulation could be a widespread phenomenon was unclear. To explore this possibility, we analysed available sequencing data of miRNA, circRNA and mRNA expression from hESC H9 cells in the undifferentiated state as well as when they have differentiated into forebrain (FB) neuron progenitor cells (45, 46). A significant proportion of circRNAs are known to be regulated along neuron differentiation –with upregulation being more frequent than downregulation (61, 62). Concomitantly, neuron-specific miRNAs are known to become more susceptible to degradation in more mature neurons (63), thus enabling a potential scenario in which circRNAs act by selectively regulating miRNA stability. In order to consider only biochemically supported circRNA-miRNA interactions, we used CLIP-Seq experimentally supported mRNA-miRNA, lncRNA-miRNA and circRNA-miRNA interaction networks catalogued in the STARBASE v3/ENCORI database (47, 48) as a proxy for bona fide interactions.

We first analyzed whether the abundance of specific miRNAs and their respective pri-miRNAs was dependent on the extent of pairing offered by circRNAs across the neuron-like differentiation process. For this analysis, we considered the variations in pri-miRNA levels as a proxy for changes in transcription rates. Next, we computed the total number of specific miRNA sites contributed by all circRNAs weighed by the average circRNA levels before and after differentiation (hereafter referred to as “miRNA-specific effective sites” on circRNAs, see materials and methods and Supplementary Dataset 1). To obtain a stoichiometrically relevant estimate of the relative circRNA pairing to which each miRNA is subjected, we further weighted the miRNA-specific effective sites to each miRNA’s average abundance before and after differentiation (ranked in Supplementary Dataset 1). Interestingly, while we observed a positive correlation between mature miRNA and pri-miRNA fold changes, the correlation became weaker for the most highly paired miRNAs (quartile “+++ paired”, Figure 5A). This allows us to suggest that while changes in mature miRNA levels can to a large extent be explained by changes in transcription of their host genes, a post-transcriptional effect might also be acting on the most highly paired miRNAs. Although we cannot rule out the possibility that circRNAs affect the processing of miRNA precursors, our analysis is consistent with a post-transcriptional effect of circRNAs on miRNA stability, acting in parallel to –but independently of– changes in miRNA transcription. This effect may occur through the concerted interaction of individual miRNAs with highly expressed circRNAs (one or multiple) across differentiation.

**Figure 5.**
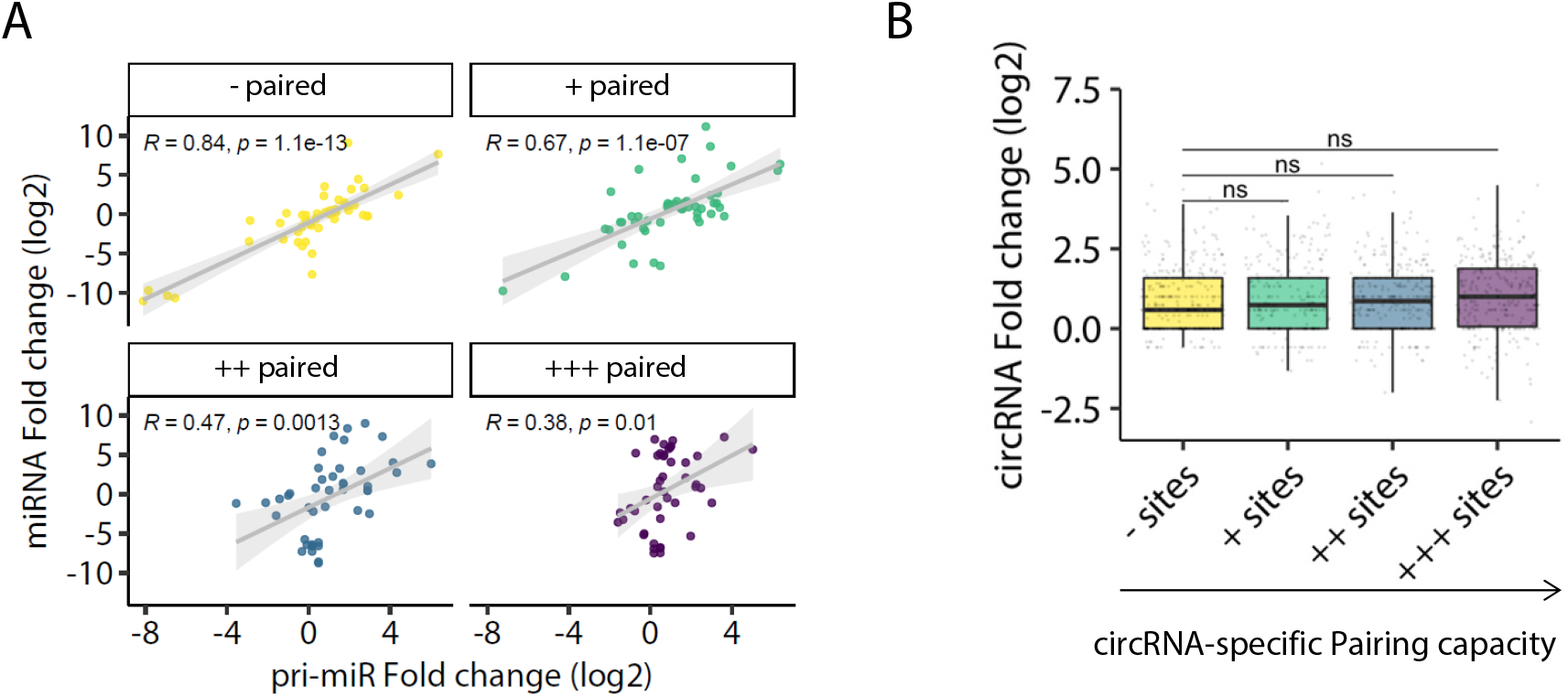
CircRNAs potentially stabilize dozens of microRNAs across neuron-like differentiation. A Scatter plot of miRNA expression fold changes (log2) across differentiation of hESC H9 cells into forebrain (FB) neuron progenitor cells plotted against the number of effective sites, coloured by quartiles of increasing number of “effective” sites within circRNAs. B Boxplot showing circRNA expression fold changes (log2) across differentiation separated by quartiles of increasing “pairing capacity” index. The analysis includes 236 miRNAs. For panel B, GLM with emmeans statistics are shown between the least paired and the remaining groups (ns: p > 0.05).

We next followed a complementary approach from a circRNA perspective, asking whether the extent of pairing offered by each circRNA, namely their ability to simultaneously interact with different miRNAs across differentiation, could impact their own abundance. To that end, we calculated a “circRNA-specific pairing capacity” index of each individual circRNA based on the total number of sites (for all miRNAs) present on the circRNA weighed by circRNA levels before differentiation (see materials and methods and Supplementary Dataset 1). We observed that an increasing “pairing capacity” index did not correlate with changes in circRNA fold changes across differentiation, which is consistent with the notion that circRNAs are not subjected to regulation by miRNAs (Figure 5B). The collection of miRNAs and circRNAs interacting across this neuron-like differentiation process includes more than a hundred miRNAs and dozens of circRNAs. In fact, by focusing on the circRNAs predicted to be most highly paired, this analysis suggests that circRNAs with the highest pairing capacity for miRNAs lie at highly connected nodes within a complex network, further supporting the view that some individual circRNAs might act as potential scaffolds for multiple miRNAs (Supplementary Figure 5A).

### Predicted binding of miRNAs to circRNAs through TDMD-like interactions

Previous knowledge indicates that a limiting number of bona fide TDMD sites –present even at sub-stoichiometric levels– can act catalytically to outcompete the tens of thousands of canonical miRNA binding sites present in the transcriptome (15, 17–19). This might seem at odds with our results indicating those TDMD sites can be either antagonized or outcompeted by a few thousand binding sites present on a subset of circRNAs. Conceivably, this conundrum could be explained if TDMD-like architectures were enriched on circRNAs themselves. In this scenario, the ability of circRNAs to resist canonical miRNA-mediated target degradation would render the TDMD-like sites present on circRNAs more effective in either competing against or triggering TDMD compared to other sites present on linear RNAs. To test this hypothesis, we searched for TDMD-like architectures within circRNAs (Figure 6A) and linear mRNAs using a TDMD-site prediction tool (55). Considering that most circRNAs are expressed from protein coding genes (64), we analysed the overlapping genomic regions (5’UTR, CDS, 3’UTR) of such genes only. Since most natural circRNAs are considered to be either not translated or only poorly translated via low efficiency cap-independent mechanisms (23), we assumed that they would be in principle largely available for miRNA pairing irrespective of the genomic region of origin due to little interference by translating ribosomes. This was further supported by the fact that we found no association between overall miRNA-pairing and the genomic region circRNAs originate from (Supplementary Figure 5D). We therefore analysed circRNAs originating from exons of all genomic regions within genes. On the other hand, for linear RNAs, we considered only 3’-UTRs within the corresponding protein-coding genes as those are the gene regions where most productive miRNA pairing occurs. Sites in 5’-UTRs and ORFs are ineffective as miRNAs appear to quickly dissociate due to displacement by the ribosome during scanning and elongation, respectively (65, 66). According to our analysis, the most highly paired miRNAs are predicted to interact through TDMD-like interactions with both linear and circRNA at a higher proportion than the least paired miRNAs, and with an overrepresented proportion of interactions on circRNAs (Figure 6B). From this analysis, we conclude that TDMD-like sites on circRNAs may be effective in affecting miRNA stability either by inducing or blocking TDMD on specific miRNAs across differentiation.

**Figure 6.**
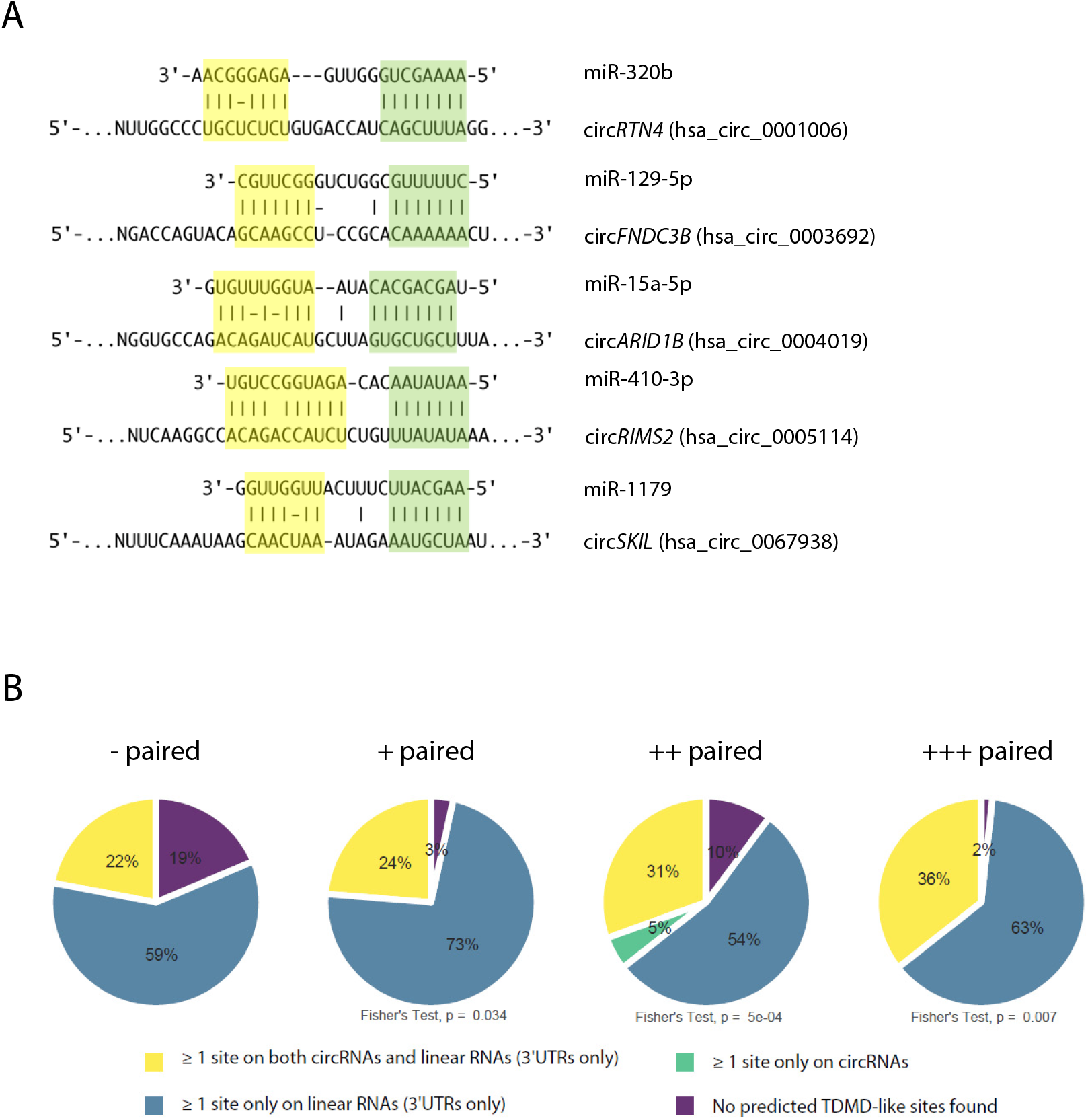
Predicted TDMD-like site architectures within circRNAs are more frequent for the most highly paired miRNAs. A Examples of predicted TDMD-like sites within circRNAs using scanMiR. All five miRNAs belong to the “+++ paired” quartile. B Pie charts showing the proportion of predicted linear RNA- and circRNA-miRNA interactions involving at least one predicted TDMD-like site against miRNAs within quartiles of increasing miRNA-specific Pairing coefficient. Shown are the p-values for the Fisher’s Exact test between each quartile and the least paired quartile (“-paired”).

## Discussion

A largely unresolved question in the field is whether circularity is intrinsically relevant for circRNAs to exert their molecular actions. Alternatively, circRNAs might have been evolutionarily selected merely based on their resistance to exonucleases that confer them special stability properties. Most experimental approaches have largely relied on knocking down or overexpressing circRNAs in different organisms or cultured cells, and undesirable artifacts can commonly act as confounding factors in the interpretation of the results. In particular, overexpression of circRNAs often suffers from a major caveat related to the co-expression of unwanted linear RNA species from commonly used expression vectors. This type of problem can easily lead to conclusions where functions are wrongly assigned to circRNAs under circumstances where the associated linear transcripts are the true functional molecules (33–35). We have attempted to address this issue by designing a strategy that allows us to express circRNAs while keeping expression of the counterpart linear product to a minimum. A similar strategy was previously reported in a different context with the goal of restricting expression of transgenes to specific tissues (67). In our hands, the approach proved to work as an efficient tool to deliver high levels of circRNAs while limiting the expression of the linear RNA derived from the circRNA construct (Figures 2, Supplementary Figure 2F).

We have exploited this tool to gain further insights into the well-known role of circRNAs in affecting miRNA function. Despite being neither prevalent nor unique to circRNAs as a class, the capacity to inhibit or “sponge” miRNA silencing activity has been the most extensively documented function of circRNAs (7, 33). In spite of this, a clear picture of the mechanism by which some circRNAs act on miRNAs is still missing. For instance, it remains unclear whether circRNAs act on miRNAs simply by blocking their function and/or by affecting their stability. It has been speculated that circRNAs might trigger TDMD similarly to some linear RNAs, but this premise was never formally and experimentally tested to date (68). Our data suggest that circular and linear RNAs carrying identical sequences differ in their TDMD properties. This is largely recapitulated by the endogenous circRNA *Cdr1as* which cannot trigger TDMD on miR-7 except when artificially expressed as a linear RNA.

The effects of *Cdr1as* on miR-7 stability might be explained by the properties of the *Cdr1as*-miR-7-Cyrano network (19). Accordingly, the stabilization of miR-7 might emerge from the inability of *Cdr1as* to drive TDMD in combination with its ability to bind and compete for miR-7, ultimately shielding the exceptionally potent TDMD activity driven by Cyrano on miR-7 (19). A similar reasoning might apply to the fact that the artificial (linear) lin*Cdr1as*, when expressed in combination with knockdown of endogenous (circular) *Cdr1as*, does not produce any additional reduction of miR-7 abundance. Under these conditions, an unleashed Cyrano-driven TDMD activity acting on miR-7 might reach a limit in miR-7 degradation rate which cannot be further enhanced by an additional TDMD-competent target such as lin*Cdr1as* (Figure 1). In contrast, in the presence of the *Cdr1as* circular RNA, the system would be set to intermediate miR-7 degradation rates which could be further enhanced by other TDMD-competent RNAs such as lin*Cdr1as* (see model in Supplementary Figure 6). Notably, a similar protection phenomenon has been observed in a prostate cancer cell line, where knockdown of circ*CSNK1G3* bearing TDMD-like binding sites for miR-181b/d decreased the abundance of these microRNAs, while circ*CSNK1G3* overexpression increased it (22) (Figure S2F).

The molecular basis explaining the functional differences between circRNAs and their cognate linear transcripts in triggering TDMD might relate to structural properties conferred by the respective circular vs. linear topologies. Previous work has shown that circRNAs possess a greater tendency to fold into more stable short local secondary structures compared to their linear counterpart transcripts (29). This may affect the flexibility of the miRNA binding region, a key factor for TDMD activity (13, 14, 20), either enhancing or limiting TDMD efficacy in different sequence contexts. In line with this idea, binding of TDMD-competent target RNAs appears to drive a conformational change on AGO proteins (loaded with specific miRNAs) that leads to their poly-ubiquitination by the ubiquitin ligase ZSWIM8 and its subsequent degradation. This would leave the specific miRNAs unprotected and susceptible to degradation by general nucleases (15, 16). The more stable folding of circRNAs could either preclude or enhance the conformational change of AGO, thus bypassing or boosting TDMD, respectively. Such folding might also limit the length-dependent effects of circRNA relative to linear RNA (Figure 4) by reducing their range of possible adopted conformations. This would be in line with our results showing that circRNA length had no influence on their impact on miRNA stability (Figure 4) or the degree of pairing to miRNAs (Supplementary Figure 5C) and with a recent report suggesting that circRNAs might bind to miRNAs more efficiently than their cognate linear RNAs (69). Alternatively, whether different target topologies lead to reduced or enhanced TDMD activity might relate to specific sequence elements that are located outside of the miRNA site *per se* for efficient triggering of TDMD (13, 16, 19, 70). Such sequences might be absent, more exposed or simply occluded within the more highly structured circRNAs, possibly explaining the differences in TDMD relative to their linear counterparts.

It might be argued that the effects of linear targets on TDMD could also relate to the presence of a 5’-cap and a poly-A tail. However, the latter seems dispensable for TDMD based on the fact that viral non-coding RNA, HSUR-1, which lacks a poly-A tail, effectively drives TDMD on miR-27 (11, 20). Likewise, by showing that circRNAs can induce TDMD, our data present the first evidence that the 5’-cap is also dispensable for this activity and that TDMD is not an exclusive property of linear RNAs.

Interactions between miRNAs and competing endogenous RNAs (ceRNAs) have received broad attention in recent years as they might represent a mechanism of miRNA inhibition (71). However, due to stoichiometry considerations, the likelihood that individual ceRNAs titrate the total amount of miRNA available for target repression seems limited (18, 72–75). Instead, models where multiple ceRNAs regulate single miRNAs have been favoured (33, 76). The case of *Cdr1as*-miR-7 pairing might represent an outstanding example that functions in an analogous way. *Cdr1as* is a highly expressed circRNA in human, rat and mouse brain with > 60 evolutionarily conserved miR-7 binding sites (1, 2), significantly exceeding the average number of sites annotated for circRNAs (based on Starbase/ENCORI database, see Supplementary Dataset 2). On the other hand, miR-7 tends to display medium to low expression, resulting in a *Cdr1as*-miR-7 stoichiometry that is compatible with *Cdr1as* protection of this miRNA. Therefore, our results favour a model where the concerted interaction of multiple circRNAs with individual miRNAs seems the most likely and relevant scenario in regulating miRNA stability. Interestingly, among the miRNAs that are upregulated upon ZSWIM8 knockdown in mouse induced neurons (15), two belong to the most highly circRNA-paired miRNAs according to our analysis (miR-7 and also miR-409-3p), suggesting that such type of regulation might be acting in neuron differentiation and possibly in pathophysiological conditions (Supplementary Figure 5B).

An increasing number of TDMD natural examples have arisen in the past few years, including both endogenous and viral transcripts (10, 11, 13, 77–80). Furthermore, the discovery that TDMD is more widespread than initially thought suggests that more examples will be discovered (15, 16, 81, 82). In this scenario, the apparent immunity of circRNAs to miRNA-mediated degradation could give them an advantage in regulating miRNA stability. This is especially true when circRNAs involve highly complementary TDMD-compatible architectures, which would lead to at least some degree of target degradation in the context of linear RNAs. Our findings support this view, suggesting that the most highly paired miRNAs seem to engage in TDMD-like interactions with circRNAs more frequently than the least paired miRNAs. Accordingly, by binding miRNAs through TDMD-like architectures, certain combinations of circRNAs might either stabilize or degrade specific miRNAs even when expressed at an overall sub-stoichiometric level relative to the whole set of linear targets present in the cell. This type of regulation could in turn be compatible with a potential reversibility of a subset of circRNA’s inhibitory function on miRNAs.

An unresolved aspect of the role of circRNAs in regulating miRNAs relates to our inability to predict their outcome on canonical miRNA silencing activity. Different outcomes may be expected depending on the binding site architectures and the relative stoichiometries of the molecules involved. For instance, miR-7 stabilization by *Cdr1as* in primary neurons and mouse cortex leads to greater average target repression (Supplementary Figure 7A-C), which is consistent with increased miRNA abundance. In contrast, miR-181b/d stabilization by circ*CSNK1G3* leads to the opposite outcome: a decreased overall target repression despite increased miRNA abundance (Supplementary Figure 7D-E). Based on this, a direct correlation between the eventual stabilization of miRNAs by circRNAs and their ensuing downstream effect on miRNA canonical silencing cannot be currently established, highlighting the need for further dissecting the role of circRNAs on miRNA target repression. Eventually, more in-depth knowledge of the players involved, their relative stoichiometries and dynamics will help us understand the emergent properties arising from different systems and the full potential and adaptive value of circRNAs in miRNA regulation.

### Limitations of the study

In this study, we have generated a set of vectors capable of expressing a circRNA with reduced levels of linear “leak” RNA that is otherwise inevitably produced from these kinds of constructs. The principle works in our hands, but the net circRNA-to-linear RNA relative expression yield that we obtain is still suboptimal for the circRNA-expressing construct bearing ZKSCAN1 introns used in neurons. Bearing this limitation in mind, our data support the notion that the circRNA levels that we achieve with this construct are enough to prevent TDMD, either by not contributing to any additional TDMD effect and/or by antagonizing the TDMD driven by its co-expressed linear counterpart. The opposite is true for the circRNA-expressing construct bearing *Lacasse2* introns tested in HEK293T cells. We are aware that both technical and biological differences might account for the observed differences.

It is important to stress that additional properties of different circRNAs might determine their ability to affect TDMD, including but not limited to circRNA modifications (RNA methylation, etc) and specific secondary structures, none of which have been systematically addressed in our work. Likewise, more studies will be required to determine if the predominant effect of circRNAs is towards blocking or enhancing TDMD.

Furthermore, we have shown that a group of miRNAs with high predicted pairing to circRNAs is potentially regulated at the post-transcriptional level across neuron-like differentiation. It is worth emphasizing that this correlation does not imply causality and that functional experiments will be needed in the future to validate this possibility. Finally, the enrichment of TDMD-like sites that we report in this study is based solely on predicted –though not experimentally validated– sites using the scanMiR bioinformatic tool (55).

## Data availability

Scripts and functions used to produce the results and plots shown in this paper can be found at: https://github.com/mmataLab/ciR-miR-stability

## Supplementary data

Supplementary Data are available at NAR online.

## Author contributions

F.F.W., D.R. and M.d.l.M. conceived the project and designed and interpreted the experiments. F.F.W. and J.L. designed, performed and interpreted most of the experiments. F.F.W. conceived, performed and interpreted most of the computational analysis. M.d.l.M. conceived, performed and interpreted some of the computational analysis. G.S. and M.S. contributed with the use of the scanMiR package to obtain the TDMD-like site enrichment results and discussed strategies for data analysis. J.L. and S.G. handled animals and prepared the neuron primary cultures. L.B. performed the subcellular fractionation experiments. F.F.W., B.P. and R.F. performed the Northern blot analysis. P.G. performed the subcloning of some of the constructs used in the study. J.P.F. designed some of the experiments and discussed experimental strategies. J.E.W., D.R. and M.d.l.M. supervised the whole project. The manuscript was written by F.F.W., D.R. and M.d.l.M.

## Acknowledgements

We thank Isabel Roditi for providing materials and discussing experiments, Helge Grosshans, Alberto R. Kornblihtt, and Javier Cáceres for a critical reading of the manuscript, Pierre-Luc Germain for a critical reading and general discussion of the manuscript, Valeria Buggiano for technical assistance and Luciana Giono for creating figure illustrations.

## Supplementary Figure Legends

**Supplementary Figure 1.**
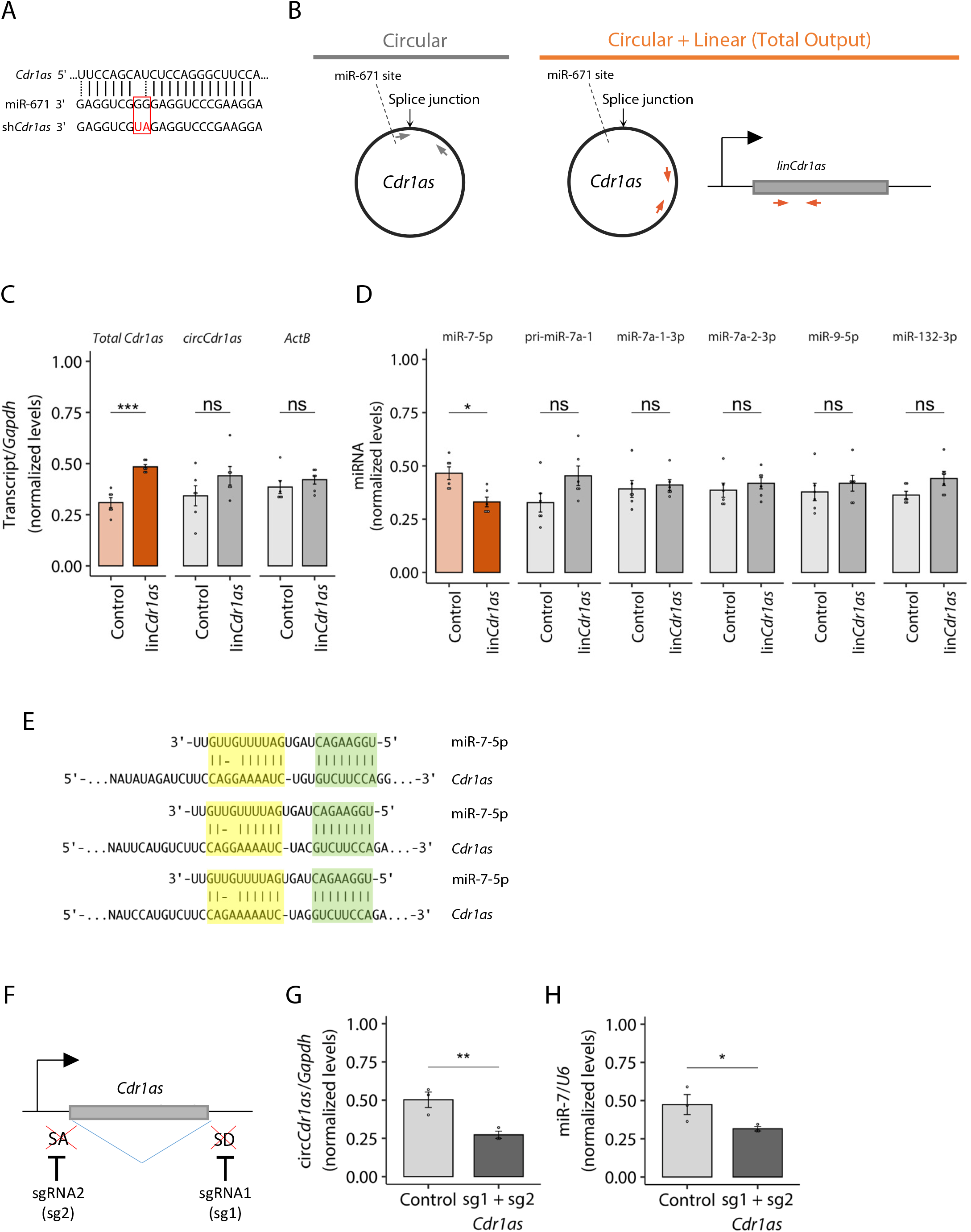
MicroRNA site illustrations, quality controls and primer design for *Cdr1*as. A. Scheme depicting the sequences of the miR-671 binding site within *Cdr1*as, miR-671 and the engineered sh*Cdr1*as. The latter is based on miR-671 with two nucleotide changes that make it fully complementary to the circRNA to maximize its slicing capacity. B. Illustration of the primer designs for measuring the different *Cdr1*as isoforms (detailed in Materials and Methods). C. Expression levels of *Cdr1*as total output, circular *Cdr1*as and unrelated gene (Actb) upon over-expression of linear *Cdr1*as (lin*Cdr1*as). Levels were normalized to Gapdh. n = 6 culture wells (from 2 independent primary cultures) for each condition. D. Expression levels of miR-7 guide strand (miR-7-5p), passenger strands (miR-7a-1-3p and miR-7a-2-3p), primary RNA (pri-miR-7a-1) and two unrelated miRNAs (miR-9-5p and miR-132-3p). All miRNAs levels were normalized to *U6*, while the pri-miR level was normalized to *Gapdh*. n = 6 culture wells (from 2 independent primary cultures) for each condition. E. Three examples of potential TDMD-competent binding sites for miR-7 present in *Cdr1*as, aligned and illustrated using scanMiR. F. Illustration of the strategy designed to mutate *Cdr1*as splicing sites by CRISPR/Cas9. G. *Cdr1*as total output levels upon CRISPR/Cas9 editing of the *Cdr1*as splicing sites, measured by RT-qPCR in primary hippocampal neurons. Control corresponds to a transduced linear transcript (GFP-expressing). n = 3 culture wells (from 1 independent primary culture) for each condition. H. MiR-7 abundance measured by Taqman RT-qPCR in the same samples as in F. n = 3 culture wells (from 1 independent primary culture) for each condition. Data are presented as mean ± SEM. Statistical significance was determined by unpaired Student’s t tests (ns: p > 0.05, *: p <= 0.05, **: p <= 0.01, ***: p <= 0.001, ****: p <= 0.0001). In panels C and D, Bonferroni’s correction for multiple testing was applied. For the F and G panels, equal variance was assumed.

**Supplementary Figure 2.**
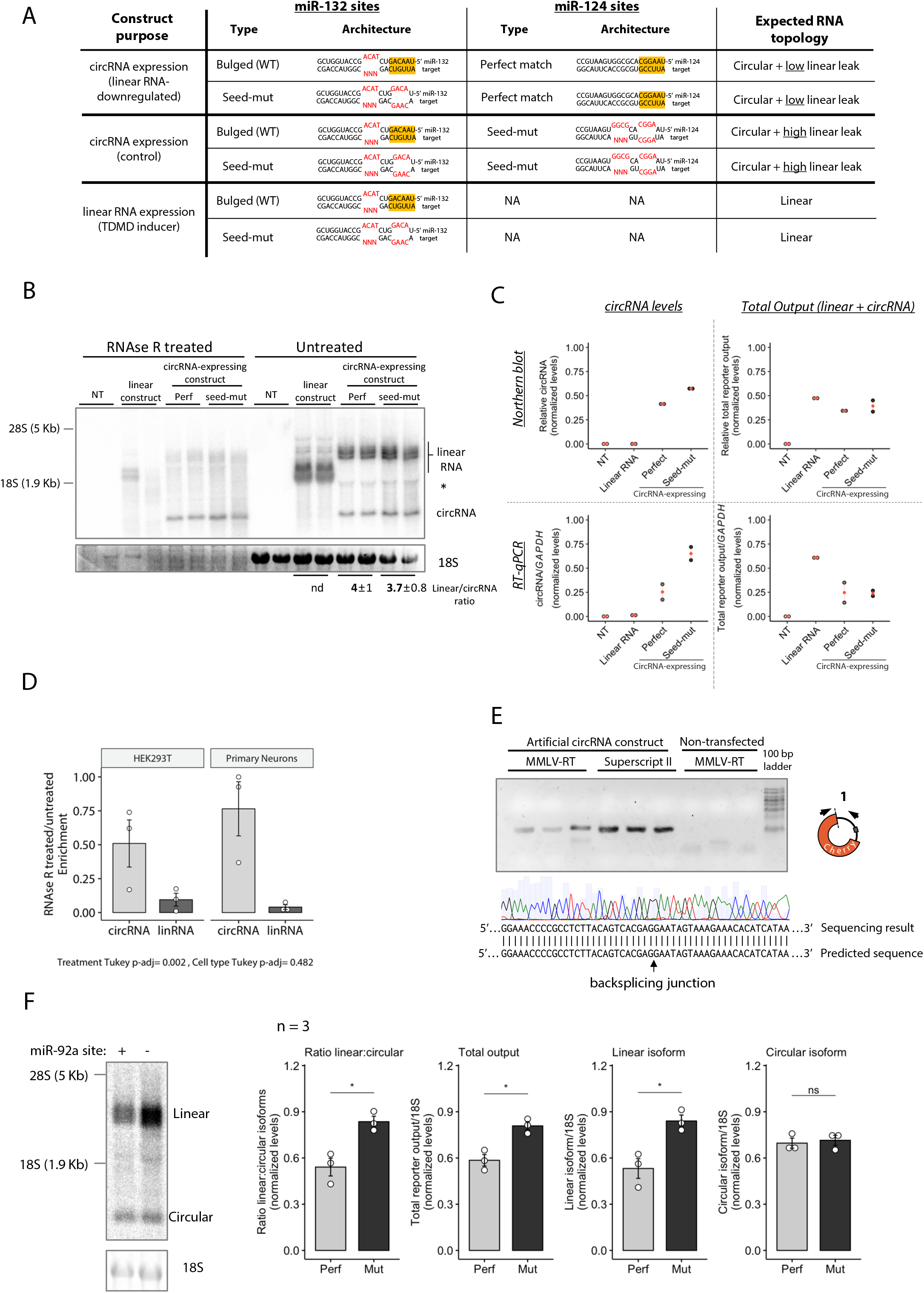
CircRNA-expressing constructs and quality controls. A. Table summarizing the generated constructs. B. Northern blot analysis of RNase R treated or untreated samples from HEK293T cells expressing the indicated constructs with either perfect or seed-mutant miR-124 binding sites. CircRNAs are resistant to RNase R digestion, confirming the circular topology of the artificial circRNA. All constructs were expressed from the tetracycline-inducible promoter (TREp). The asterisk (*) marks a putative mechanically linearized product of the corresponding circRNA. Linear/circRNA ratio quantification was performed by digital densitometry using ImageJ. C. Quantification by different methods of the linear and circular isoforms from RNase R untreated samples of HEK293T cells (naturally lacking miR124) expressing the indicated constructs. Top panels: digital densitometry quantification of Northern blot bands by ImageJ. Bottom panels: RT-qPCR quantification using primers specific for the circular isoform or for both the linear and circular isoforms combined (Total output) as depicted in Figure 2A. In red, the mean is indicated. n = 2 culture wells (from 1 experiment) of HEK293T cells for each condition. D. RT-qPCR measurement of the circRNA-to-linear RNA ratio of RNase R treated vs. untreated samples in cortical neurons or HEK293T cells using divergent primers depicted in Figure 2. Values reflect relative rather than absolute ratios of the measured isoforms. n = 3 culture wells (from 1 primary culture/experiment) of each cell type and for each condition. E. Top: Agarose gel showing triplicates of the RT-qPCR amplicons obtained with divergent primers against the backsplicing junction of the artificial circRNA, after retro-transcription with two different reverse transcriptases (MMLV-RT & Superscript II) in order to rule out artifacts due to template switching during cDNA synthesis. Bottom: Sanger sequencing of the amplicons shown above confirming backsplicing junction in HEK293 cells. F. Left: Northern blot analysis of in HEK293T cells expressing the indicated constructs bearing either perfect or seed-mutant miR-92a binding sites. Right: digital densitometry quantification of Northern blot bands by ImageJ. In (F), data are presented as mean ± SEM. Statistical significance was determined by unpaired Student’s *t* tests (ns: p > 0.05, *: p <= 0.05, **: p <= 0.01, ***: p <= 0.001, ****: p <= 0.0001).

**Supplementary Figure 3.**
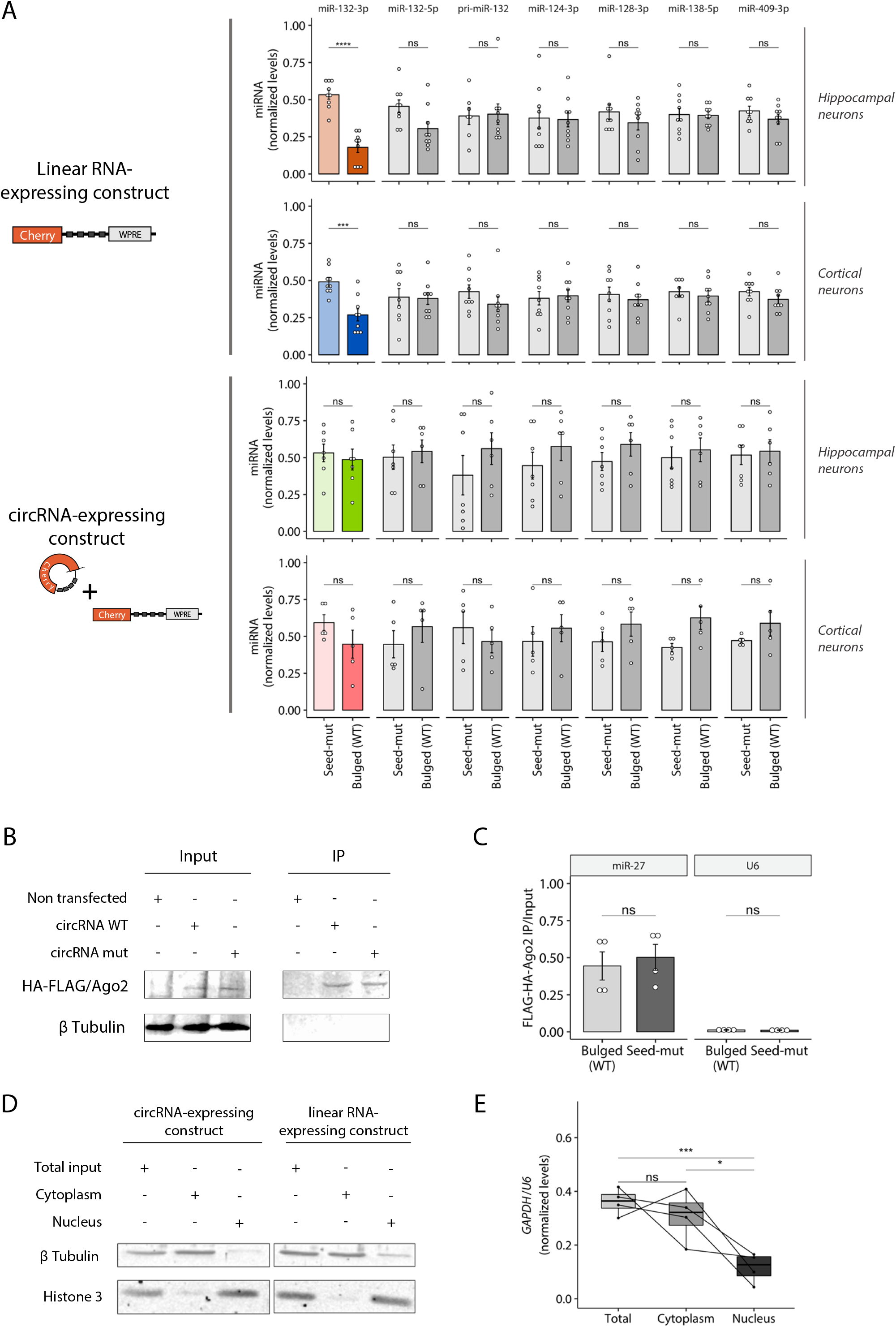
TDMD specificity, HA-FLAG/AGO2 RIP and cellular fractionation quality controls. A. RT-qPCR quantification of the indicated RNA species confirms specific degradation of miR-132 guide strand (miR-132-3p) and discards potential transcriptional effects. Levels of miR-132 passenger strand (miR-132-5p), primary transcript (pri-miR-132) and four unrelated miRNAs (miR-124-3p, miR-128-3p, miR-138-5p and miR-409-3p) were normalized to *U6*. n = 9 culture wells (from 3 independent primary cultures) of cortical neurons for each condition; n = 9 culture wells (from 4 independent primary cultures) of hippocampal neurons for each condition. Missing points are failed culture wells/RT-qPCR reactions. B. Representative anti-HA Western Blot from inputs or anti-FLAG IPs in non-transfected HEK-293T cells or cells co-transfected with HA-FLAG/AGO2 and the circRNA-expressing construct bearing bulged (WT) or seed-mutant (mut) miR-132 sites. C. *U6* and miR-27 levels were measured to verify the efficiency of HA-FLAG/AGO2 immunoprecipitation. n = 4 culture wells (from 2 experiments) of HEK293T cells for each condition. D. Representative Western Blot of HEK293T cells transfected with either the circRNA-expressing construct or the linear control RNA, following subcellular fractionation. β-Tubulin and Histone-3 were used as cytoplasm and nucleus markers, respectively. E. Boxplot depicting median values of *GAPDH/U6* ratio measured by RT-qPCR confirms proper subcellular fractionation of samples either transfected with the circRNA-expressing construct or the linear control. n = 4 culture wells (from 1 experiment) of HEK293T cells for each condition. In (A, C), data are presented as mean ± SEM. Statistical significance was determined by unpaired Student’s *t* tests (ns: p > 0.05, *: p <= 0.05, **: p <= 0.01, ***: p <= 0.001, ****: p <= 0.0001). In panel A, Bonferroni’s correction for multiple testing was applied.

**Supplementary Figure 4.**
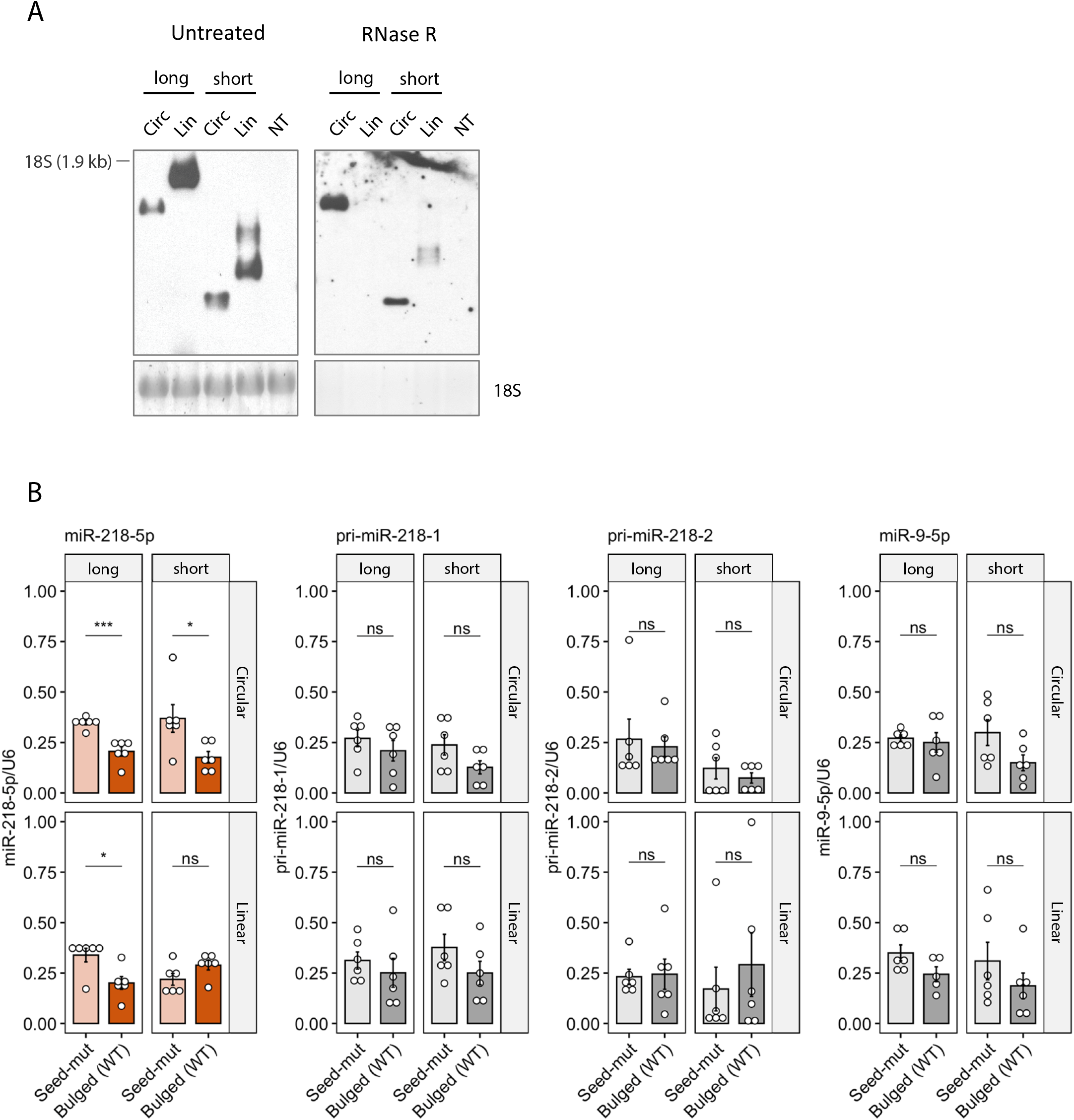
CircRNA-expressing constructs and quality controls. A. Northern blot analysis of RNase R treated or untreated samples from HEK293T cells expressing the constructs of the indicated length. CircRNAs are resistant to RNase R digestion, confirming the circular topology of the artificial circRNA. B. RT-qPCR quantification of the indicated RNA species confirms specific degradation of miR-218 guide strand (miR-218-5p) and discards potential transcriptional effects. Levels of primary transcript (pri-miR-218) and one unrelated miRNA (miR-9-5p) were normalized to *U6*. n = 6 culture wells (from 2 independent experiments) of HEK293T cells for each condition. Data are presented as mean ± SEM. Statistical significance was determined by unpaired Student’s t tests (ns: p > 0.05, *: p <= 0.05, **: p <= 0.01, ***: p <= 0.001, ****: p <= 0.0001).

**Supplementary Figure 5.**
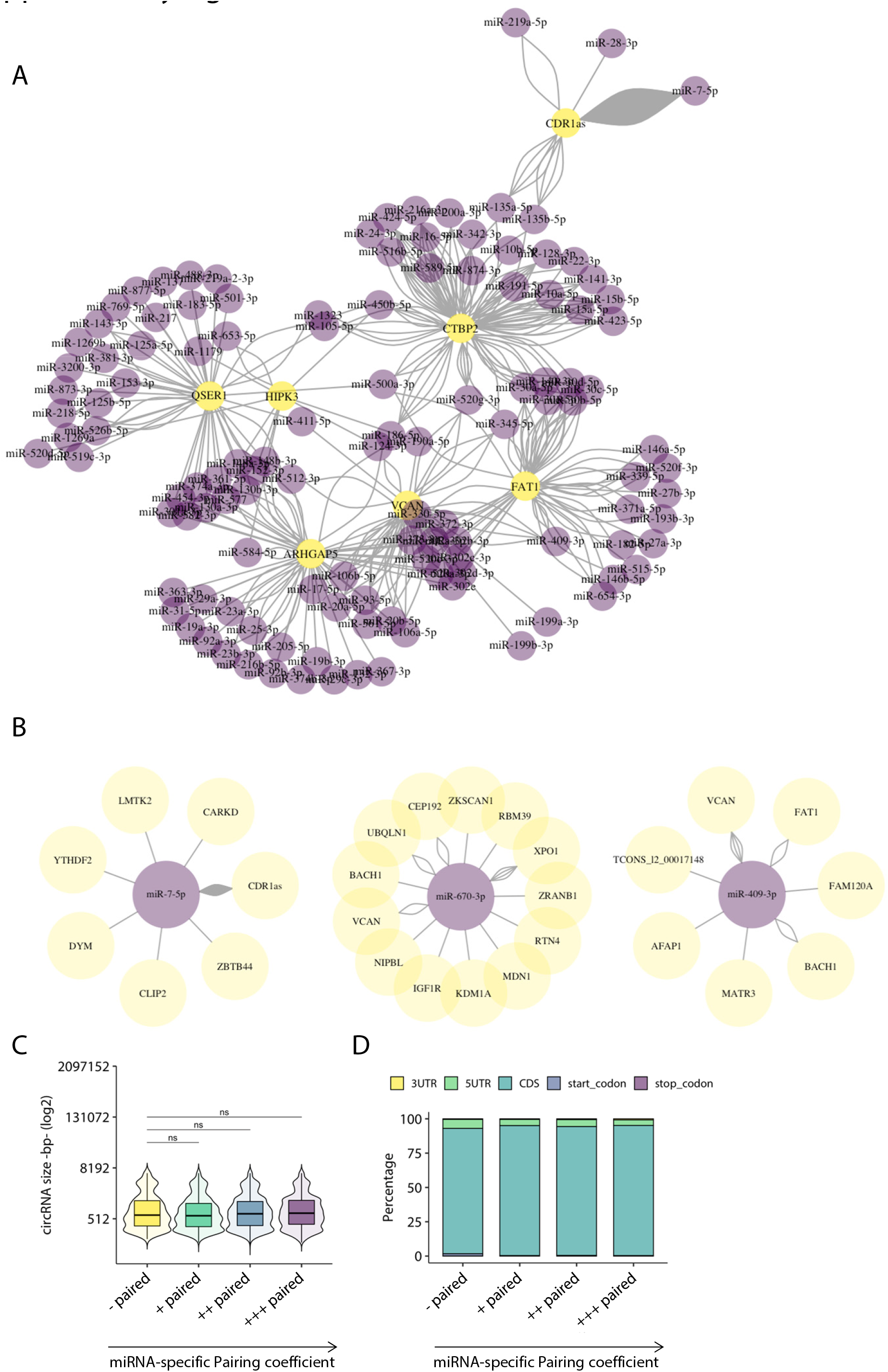
Illustrations of the predicted circRNA-miRNA networks and genomic features of the potentially involved circRNAs. A. Network diagram of biochemically supported interactions between circRNAs and miRNAs. Depicted are the interactions involving the seven most “pairing” circRNAs. B. Network diagram showing biochemically supported circRNA-miRNA interactions for miR-7 and two other miRNAs with similar “pairing” coefficient (miR-670-3p and miR-409-3p). For all three examples, the stoichiometries (i.e. ratios of circRNA-binding-site:miRNA) are similar, with *CDR1*as making the largest contribution for miRNA-7. MiR-7 and miR-409 are also a demonstrated case and strong candidate to undergo of TDMD, respectively (see main text). C. Boxplot showing the size of circRNAs (log2 of the base pairs) that interact with miRNAs from each of the different pairing quartiles estimated from data of hESC H9 into forebrain (FB) neuron progenitor cell differentiation (Chen *et al*, 2015; Zhang *et al*, 2016). Shown are Wilcoxon rank sum test p-values (corrected with the Hochberg method for multiple comparisons) between the least paired and the remaining groups (ns: p > 0.05). D. Barplot representing the percentages of overlapping genomic regions (CDS, 5’UTR, 3’UTR, etc) giving rise to the circRNAs with predicted binding sites against miRNAs within different quartiles of miRNA-specific Pairing coefficient. No enrichment of genomics features for circRNAs interacting with miRNAs within the different quartiles was observed (Pearson’s Chi-squared test p-value = 0.998).

**Supplementary Figure 6.**
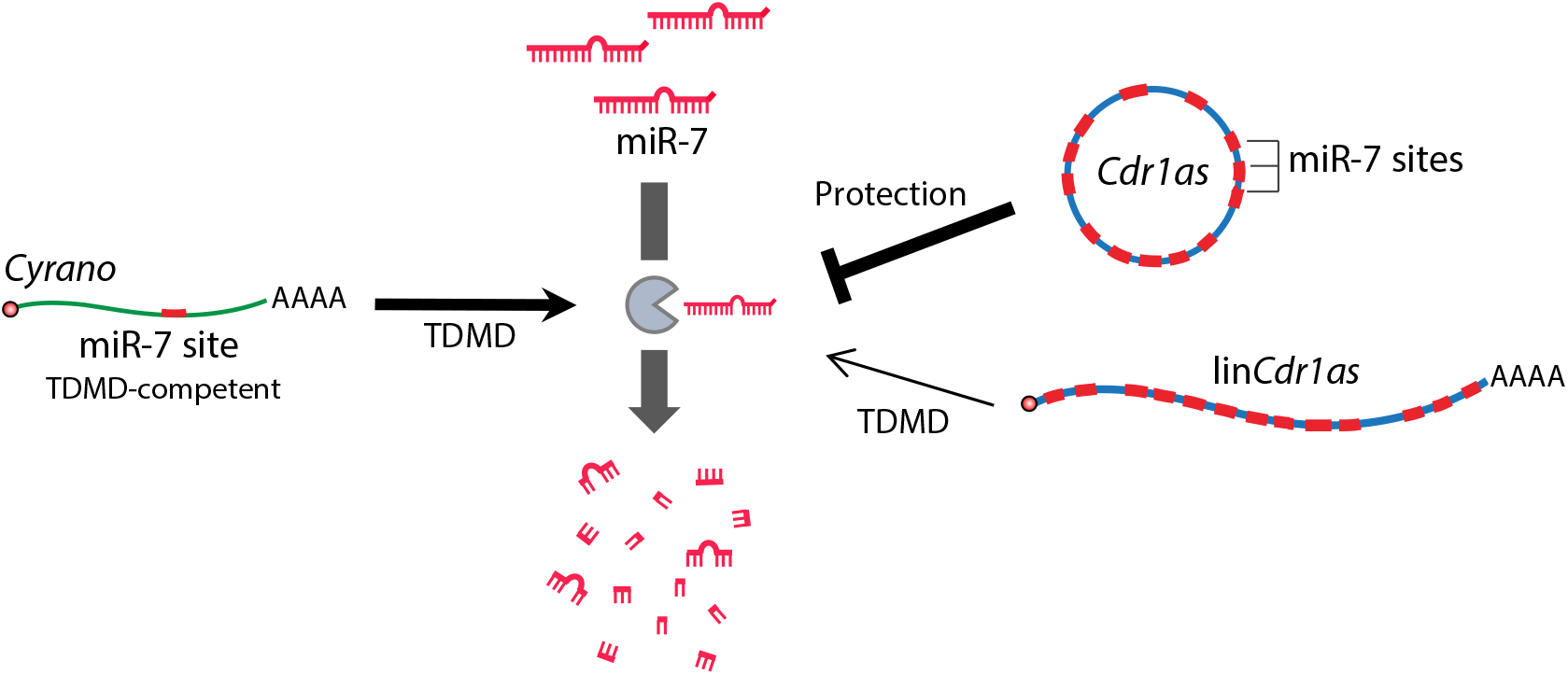
Model for the topology-dependent effect of *Cdr1*as on miR-7.

**Supplementary Figure 7.**
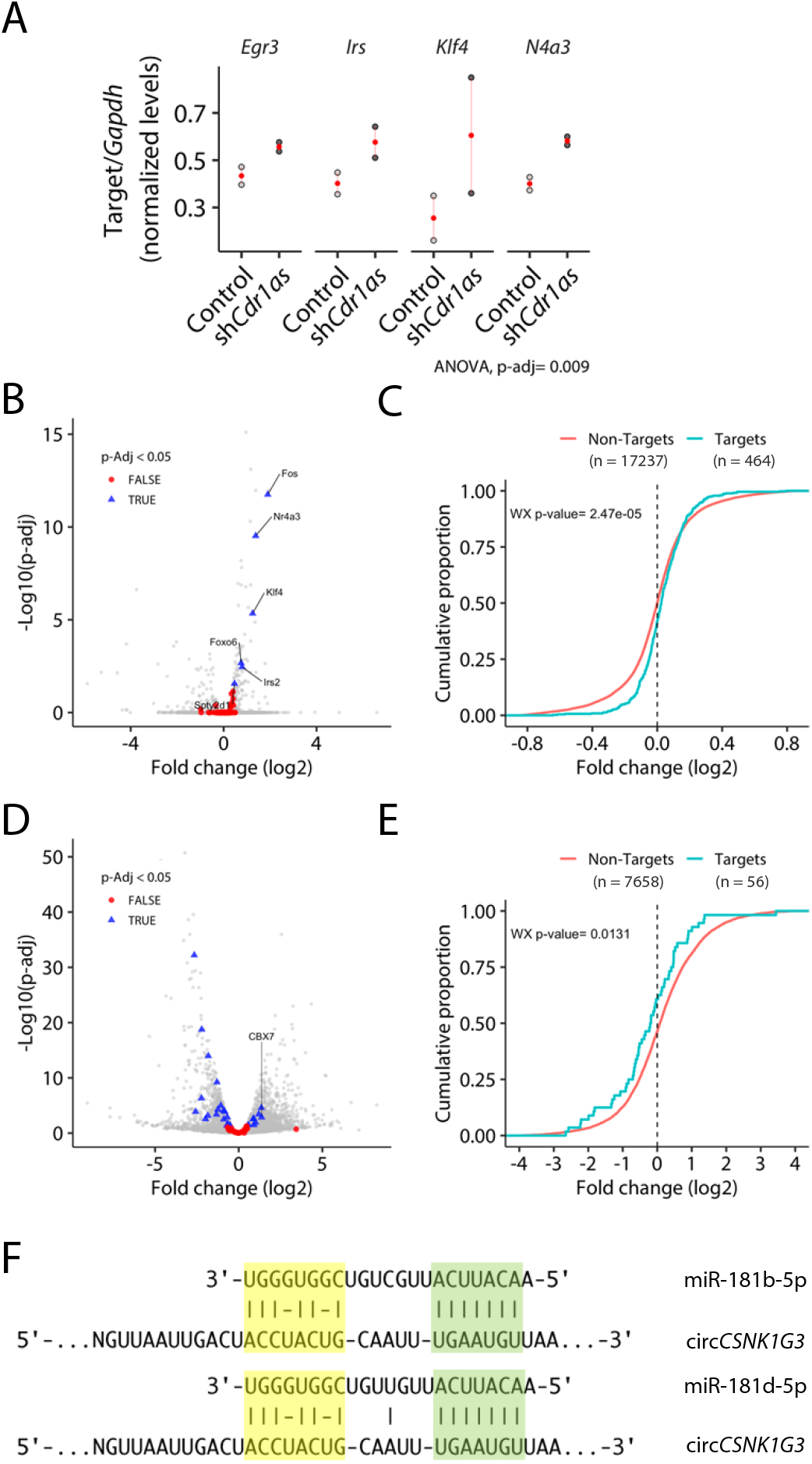
MiRNA stabilization by circRNAs leads to contradictory effects on the overall target silencing in different systems. A Abundance of four validated miR-7 targets, measured by RT-qPCR upon Cdr1as knockdown in cortical primary neurons. As previously reported, all four targets were upregulated upon *CDR1*as KO relative to control (Piwecka et al, 2017). In red, the mean and its standard error (SEM). B-C Re-analysis of data from Piwecka et al. 2017 confirming that miR-7 predicted targets are significantly upregulated in mouse cortex upon *CDR1*as knockout. (B) Volcano plot depicting fold-changes (log2) of miR-7 predicted targets in red (p-Adj > 0.05) or blue (p-Adj < 0.05) and non-targets in grey. (C) Fold change (log2) distribution of predicted miR-7-5p targets compared to the background (Non-Targets) in cortex of *Cdr1*as KO vs. WT mice. Wilcoxon–Mann–Whitney (WX) test p-values are shown. D-E Re-analysis of data from Chen et al. 2019 to measure average effects on miR-181b/d predicted targets in circCSNK1G3 KD vs. WT PC-3 prostate cancer cells. (D) Volcano plot depicting fold changes (log2) of miR-181b/d predicted targets in red (p-Adj > 0.05) or blue (p-Adj < 0.05) and non-targets in grey. Note that in opposition to the general trend, CBX7 abundance is upregulated upon circCSNK1G3 knockdown, which recapitulates published results (Chen et al, 2019). (E) Fold change (log2) distribution of predicted miR-181b/d targets compared to the background (Non-Targets) in circCSNK1G3 KD vs. WT PC-3 prostate cancer cells. Wilcoxon–Mann–Whitney (WX) test p-values are shown. F ScanMiR alignment of the miR-181b/d predicted sites on circCSNK1G3 shows extensive 3’ end complementarity. Fold changes (log2) and p-Adj values were calculated using the DESeq2 package in R.

